# Novel reassortments of variation underlie repeated adaptation to whole genome duplication in two outcrossing Arabidopsis species

**DOI:** 10.1101/2023.01.11.523565

**Authors:** Magdalena Bohutínská, Eliška Petříková, Tom R. Booker, Cristina Vives Cobo, Jakub Vlček, Gabriela Šrámková, Alžběta Poštulková, Jakub Hojka, Karol Marhold, Levi Yant, Filip Kolář, Roswitha Schmickl

## Abstract

Polyploidy, the result of whole genome duplication (WGD), is widespread across the tree of life and is often associated with speciation or adaptability. It is thought that adaptation in autopolyploids (within-species polyploids) may be facilitated by increased access to genetic variation. This variation may be sourced from gene flow with sister diploids and new access to other tetraploid lineages, as well as from increased mutational targets provided by doubled DNA content. Here we deconstruct the origins of haplotype blocks displaying the strongest selection signals in established, successful autopolyploids, *Arabidopsis lyrata* and *Arabidopsis arenosa*. We see strong signatures of selection in 17 genes implied in meiosis, cell cycle, and transcription across all four autotetraploid lineages present in our expanded sampling of 983 sequenced genomes. Most prominent in our results is the finding that the tetraploid-characteristic haplotype blocks with the most robust signals of selection were completely absent in diploid sisters. In contrast, the fine-scaled variant mosaics in the tetraploids originated from highly diverse evolutionary sources. These include novel reassortments of trans-specific polymorphism from diploids, new mutations, and tetraploid-specific inter-species hybridization. We speculate that this broad-scale allele acquisition and re-shuffling enabled the autotetraploids to rapidly adapt to the challenges inherent to WGD, and may further promote their adaptation to environmental challenges.

**Lay summary:** Polyploidy, the result of whole genome duplication, is associated with speciation and adaptation. To fuel their often remarkable adaptations, polyploids may access and maintain adaptive alleles more readily than diploids. Here we identify repeated signals of selection on genes that are thought to mediate adaptation to whole genome duplication in two *Arabidopsis* species. We found that the tetraploid-characteristic haplotype blocks, found in genes exhibiting the most robust signals of selection, were never present in their diploid relatives. Instead, these blocks were made of mosaics forged from multiple allelic sources. We hypothesize that this increased variation helped polyploids to adapt to the process that caused this increase – genome duplication – and may also help them adapt to novel environments.

## Introduction

Whole genome duplication (WGD) is widespread across eukaryotes, especially in plants. It comes with significant costs, such as meiotic instability and cell cycle changes, both of which require adaptation (Bomblies, 2020; Doyle & Coate, 2019). Over the last decade, a body of work has accrued describing means by which within-species WGD lineages (autopolyploids) overcome these challenges. The most obvious of these appears to be a stabilization of cell division via selection acting at meiotic and mitotic genes (Bohutínská, Alston, et al., 2021; Bohutínská, Handrick, et al., 2021; Bray et al., 2023; Yant et al., 2013). Additionally, cyclin genes have been observed to mediate tolerance to tetraploidization in polyploid tumours and in proliferating *Arabidopsis* tissues (Crockford et al., 2017; Imai et al., 2006; Potapova et al., 2016; Sterken et al., 2012). And while certain genes involved in adaptation to WGD have been identified and their functions verified (Morgan et al., 2022; Morgan et al., 2020), a broader evolutionary context remains elusive. Questions persist regarding the origins of adaptive genetic variation and the mechanisms by which these variants assemble into positively selected haplotype blocks in the nascent polyploids.

From a genetic point of view, polyploids may benefit from enhancement of particular traits that facilitate adaptation. In diploids, various sources contribute to the pool of adaptive alleles, including gene flow/introgression, de novo mutations, and ancestral standing variation (Fig. 1A). Recent work indicates the potential for all these sources to become greater following WGD (Fig. 1B). First, polyploid lineages can benefit from the removal of hybridization barriers (Lafon-Placette et al., 2017; Marhold & Lihová, 2006; Schmickl & Yant, 2021), leading to increased allelic exchange through introgression (Arnold et al., 2016; Marburger et al., 2019). Polyploids can also gain additional diversity via introgression from sister diploids, whereas introgression in the opposite direction is much less likely due to the nature of the diploid-tetraploid barrier (Baduel et al., 2018; Morgan et al., 2021; Te Beest et al., 2012). Second, polyploids experience elevated mutational input due to the doubled chromosome numbers, leading to a higher number of novel mutations per generation (Selmecki et al., 2015). Finally, polyploids maintain genetic diversity more efficiently due to increased masking of recessive alleles (Monnahan et al., 2019; Ronfort, 1999), thus keeping a larger pool of standing variation (Hämälä et al., 2023). Consequently, autopolyploids generate and maintain a greater degree of genetic variability than diploids (Fig. 1).

**Fig. 1:**
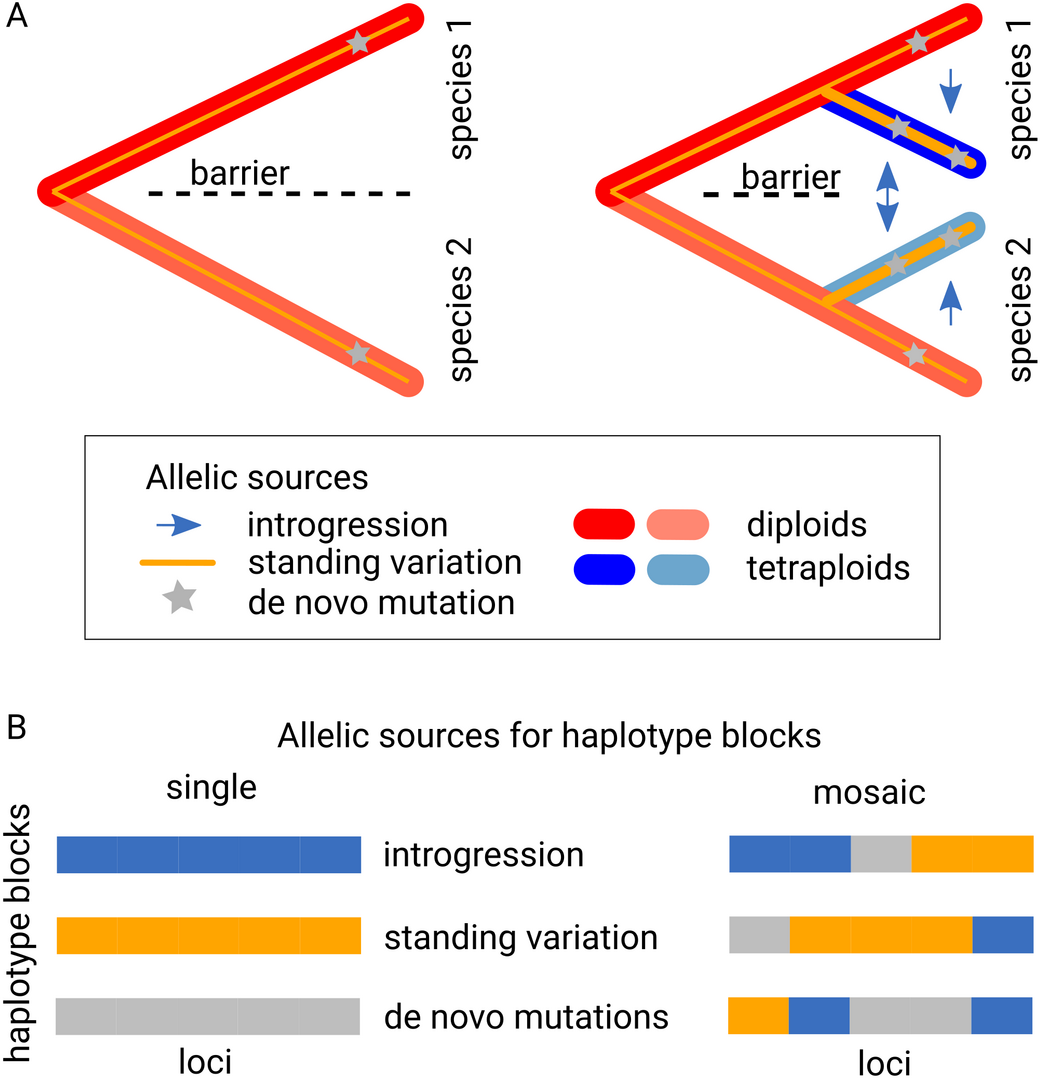
Hypotheses about allelic sources in a diploid-autotetraploid system. A: As compared to diploids, autotetraploid lineages may acquire alleles via increased introgression potential, increased population-scaled mutational input, and higher level of retainment of ancestral standing variation. Shown are two diploid species (red), each of which gave rise to an autotetraploid (blue) lineage (right). B: Higher variability in the possible sources of potentially beneficial alleles suggests that positively selected haplotype blocks in tetraploids might be more likely to form from a mosaic of sources (right), in contrast to the typically assumed homogeneous (single-source) scenario (left).

Beneficial alleles are commonly envisioned as originating from a single source (as shown in Fig. 1), either from introgression, standing genetic variation, or de novo mutations (Lee & Coop, 2019, Bohutínská, Vlček, et al., 2021; Konečná et al., 2021; Oziolor et al., 2019; Wang et al., 2021). However, recent evidence suggests that alleles can also gradually accrue to form finely tuned haplotypes (Archambeault et al., 2020; Roberts Kingman et al., 2021; Seear et al., 2020). This implies that adaptive haplotypes may accumulate from multiple sources, rather than just one. Here we ask if polyploids use their expanded allelic sources to construct novel fine-scaled mosaic haplotype blocks of diverse origins, and what those origins are.

The naturally ploidy-variable (diploid and autotetraploid) species *Arabidopsis arenosa* and *Arabidopsis lyrata* have emerged as powerful models to understand adaptation to WGD (Yant & Bomblies, 2017). Recent research in these species identified shared candidate alleles mediating adaptation to WGD (Bohutínská, Handrick, et al., 2021; Marburger et al., 2019; Seear et al., 2020; Yant et al., 2013). However there has been no comprehensive investigation of the evolutionary origin of such tetraploidy-related alleles across broad sampling of the species, lineages, and loci involved. There have been indications that candidate adaptive alleles might have originated from specific source populations, or that recombination may be involved in the construction of adaptive alleles (Marburger et al., 2019, Seear et al., 2020). However, these studies were limited by small sample sizes that covered only a fraction of each species range. Additionally, they did not consider trans-specific polymorphism (here represented by ancestral alleles shared among different *Arabidopsis* species). We thus do not know to which extent the different allelic sources contributed to the formation of potentially adaptive tetraploid haplotype blocks.

Here we take advantage of an exhaustive dataset of 983 sequenced individuals encompassing all known lineages of European *Arabidopsis* autotetraploids, backed by genome-wide diversity of all diploid outcrossing *Arabidopsis* species. Apart from the tetraploid *A. lyrata* lineage from the eastern Austrian Forealps (‘Austria’ hereafter; (Marburger et al., 2019; Schmickl & Koch, 2011)), we newly analysed two additional *A. lyrata* tetraploid lineages from Central Europe: south-eastern Czechia (‘Czechia’ hereafter) and Harz in Germany (‘Germany’ hereafter). We performed a joint analysis of these three *A. lyrata* tetraploid lineages and tetraploid *A. arenosa*, which derived from a single WGD event (Arnold et al., 2015, Monnahan et al., 2019). We used this sampling of four tetraploid lineages, established which genes and processes are robustly under selection in each lineage, and elucidated the sources of candidate adaptive alleles. Using this knowledge, we asked to which extent tetraploid-specific haplotype blocks were formed and what were their sources.

## Methods

### Sampling

*Arabidopsis*, a well-studied plant model genus, is primarily diploid, but autotetraploids have been discovered among the diverse outcrossing species *A. arenosa* and *A. lyrata* in Central Europe. While *A. arenosa* diploids and tetraploids are widespread, *A. lyrata* primarily occupies a narrow ecological niche in Central Europe. Previous research suggested that introgression between *A. arenosa* and *A. lyrata* tetraploids facilitated the genetic differentiation of *A. lyrata* tetraploids from their diploid ancestors, allowing them to expand their niche (Marburger et al., 2019; Schmickl & Koch, 2011).

Here we sampled and sequenced genomes from both diploid and neighboring tetraploid populations across all known diploid-tetraploid lineages in Central European *Arabidopsis*. Our dataset includes genomes from total 73 newly sequenced diploid and tetraploid individuals of both species. Additionally, we incorporated data from 910 previously published whole genome sequences (Bohutínská, Vlček, et al., 2021; Guggisberg et al., 2018; Hämälä et al., 2017; Hämälä et al., 2018; Konečná et al., 2021; Marburger et al., 2019; Mattila, et al., 2017; Monnahan et al., 2019; Novikova et al., 2016, 2017; Preite et al., 2019), encompassing a total of 983 individuals and 129 populations (Dataset S1, Dataset S7). The ploidy level of each individual in our new sampling was confirmed using flow cytometry following (Kolář, Lučanová, et al., 2016).

### Population genetic structure

Samples were whole genome re-sequenced (as detailed in Dataset S1), and single nucleotide polymorphisms (SNPs) were called using a ploidy-aware approach following (Bohutínská, Vlček, et al., 2021; Monnahan et al., 2019). *Arabidopsis arenosa* and *A. lyrata* autotetraploids show random segregation of chromosomes (Arnold et al., 2015; Seear et al., 2020), resulting in allele frequency estimation comparable to diploids (Meirmans et al., 2018). This allowed to use a set of methods designed for diploids, which are also applicable to mixed-ploidy data; for a discussion see (Bohutínská et al., 2023)). Specifically, we inferred relationships between populations using allele frequency covariance graphs implemented in TreeMix v1.13 (Pickrell & Pritchard, 2012). *Arabidopsis arenosa* was rooted with the diploid Pannonian population BDO and *A. lyrata* with the diploid Scandinavian population LOM. To obtain confidence in the reconstructed topology of *A. arenosa* and *A. lyrata* (Fig. 1B, C), we performed a bootstrap analysis with 1000 bp blocks (matching the selection scan window size below) and 100 replicates. Further, we used Bayesian clustering in FastStructure (Raj et al., 2014). We randomly sampled two alleles per tetraploid individual using a custom script. This approach does not appear to bias clustering in autotetraploid samples based on (Stift et al., 2019). Finally, we displayed genetic relatedness among individuals using principal component analysis (PCA) as implemented in adegenet (Jombart, 2008). We calculated genome-wide four-fold degenerate (4d) within-population metrics (nucleotide diversity (π) and Tajima’s D (Tajima, 1989)) using the python3 ScanTools_ProtEvol pipeline (Bohutínská, Vlček, et al., 2021).

To test for introgression, we used the ABBA-BABA test as described in (Martin et al.,2013). We assessed introgression from tetraploid *A. arenosa* to each *A. lyrata* tetraploid lineage. For the non-admixed population (P1), we used the early diverging diploid Pannonian lineage of *A. arenosa* (Kolář, Fuxová, et al., 2016), population BDO. Allele frequencies were polarised using diploid *A. halleri*, population GUN from Austria.

### Genome-wide scans for positive selection

To find candidate positively selected genes (PSGs) which likely contributed to adaptation to whole genome duplication in tetraploids, we employed two selection scan methods suitable for both diploids and autotetraploids (Bohutínská et al., 2023; Booker et al., 2023).

First, between each of the tetraploid populations and the most proximal diploid population, we calculated Fst (Hudson, Slatkin, & Maddison, 1992) in 1 kb windows. We then used PicMin to test whether there was evidence of repeated genetic differentiation among the tetraploid/diploid population pairs. PicMin uses order statistics to test whether population genetic summary statistics (in this case Fst) for orthologous genomic regions in different lineages exhibit a common shift towards extreme values in multiple species, indicative of repeated action of positive selection. PicMin was applied on windows that had data for at least three lineages (86,249 windows in total). A genome-wide false discovery rate correction was then performed with a significance threshold of q < 0.01. In cases of outlier signal spanning adjacent windows, the window with the lowest q-value and highest Fst was retained.

We also employed a ‘candidate SNP’ approach by calculating SNP-based Fst between tetraploid and their most proximal diploid populations within each lineage. A 1% outlier threshold was used to identify highly differentiated SNPs (‘candidate SNPs’ hereafter) across the genome. We determined the density of these candidate SNPs per gene using *A. lyrata* gene models (Rawat et al., 2015). Candidate genes were identified as the upper quartile with the highest density of outlier SNPs. Fisher’s exact test (‘SuperExactTest’ package (Wang et al., 2015) in R) was then applied to identify repeatedly identified candidates. Notably, all PicMin-identified genes were confirmed by this candidate SNP approach.

### Functional annotation

To infer functions significantly enriched in our list of tetraploid PSGs, we performed gene ontology (GO) and UniProt Keywords enrichment analyses using the STRING database (https://string-db.org/, last accessed 02/03/2023, (Szklarczyk et al., 2015)). We used *Arabidopsis thaliana* orthologs of *A. lyrata* genes. Only categories with FDR < 0.05 were considered. We also manually searched for functional descriptions of each gene using the TAIR database and associated literature (Berardini et al., 2015). To identify protein-protein interactions, we used the STRING database, including all available information sources, and focused on 1st shell interactions (Szklarczyk et al., 2015).

### Allele frequencies of haplotype blocks

We reconstructed haplotype blocks for the 12 PSGs (a subset of 17 PSGs supported by at least five ‘candidate SNPs’) within *A. lyrata* and *A. arenosa* populations based on linked marker SNPs (232 ‘candidate SNPs’ in total; Dataset S4), following a procedure by (Bohutínská, Handrick, et al., 2021). Haplotype block frequencies were calculated separately for diploids (218 *A. arenosa* and 121 *A. lyrata* individuals) and tetraploids (479 individuals). For each of the 12 PSGs with *n* candidate SNPs each (for *n* see Fig. 3B), we determined the allele frequency (*AF*) of major allele across all individuals. With 1916 tetraploid haplotypes and 436 (*A. arenosa*) / 242 (*A. lyrata*) diploid haplotypes, the major *AF* corresponds to the tetraploid allele. We defined the tetraploid haplotype block frequency as the minimum among *n AF*s (HAFt = min {major *AF*}), and the *AF* of diploid haplotype blocks as HAFd = 1 – max {major *AF*}. The *AF* of all other haplotype blocks resulting from recombination or new mutations was defined as HAFr = 1 – HAFt – HAFd. Calculations were performed using an in-house R script.

To infer allele frequencies of haplotype blocks, we used a SNP frequency-based approach due to unreliable standard phasing algorithms in short read tetraploid samples (Kyriakidou et al. 2018). We thus validated our haplotype block analysis by estimating the correspondence between our allele frequency-based inferred haplotypes and (1) sequences from PacBio HiFi-based long read assemblies of the 12 PSGs in five diploid and five tetraploid samples of *A. arenosa*, and (2) sequences from long reads themselves in two PSGs, two diploid and two tetraploid samples. For (1), we used BLASTn v2.10.0 to extract sequences from the 12 PSGs in newly produced diploid and tetraploid assemblies (Dataset S5). These sequences were then aligned with MUSCLE and visualized with IGV v2.11.9 (Dataset S4). For (2), we aligned the long reads spanning the region of PSGs CYCA2;3 and PDS5b to the assemblies, visualized with IGV, and manually exported the information about physical linkage of candidate SNPs into (Dataset S6).

### Allelic sources of tetraploid haplotype blocks

To assess if tetraploid haplotype blocks in our study originated from a single source or as a mosaic of allelic sources, we employed two methods: tracing the evolutionary origins of candidate SNPs across the ancestral diploid lineages (Novikova et al., 2016), and reconstructing networks of genetic distances at the PSG regions (Jombart, 2008).

To determine the allelic sources contributing to tetraploid haplotype blocks, we investigated the occurrence of variants defining the tetraploid haplotype blocks (the 232 ‘candidate SNPs’) within diploid individuals. We used a comprehensive genomic dataset encompassing diploid *A. lyrata* and *A. arenosa* (339 diploid individuals) alongside their outcrossing relatives *A. halleri, A. croatica, A. cebennensis*, and *A. pedemontana* (‘diploid congeners’, totaling 165 individuals; see Dataset S1). A rarefaction analysis in (Bohutínská, Handrick, et al., 2021) indicated that sampling as few as 40 diploid individuals across the *A. arenosa* species range captures the majority of diploid diversity. Therefore, our dataset of 504 diploid individuals should suffice to cover the full natural diversity of diploids.

We excluded singletons (variants occurring only once in a species) to reduce the impact of sequencing errors (results were robust with or without singletons; Fig. S6). Then, using species’ phylogenetic relationships (Fig. 5A, (Novikova et al., 2018)), we determined likely allelic sources for the 232 candidate SNPs among 504 diploid individuals. We categorized each SNP according to one of the following seven scenarios (Fig. 5A): 1) trans-specific polymorphism in both *A. lyrata* and *A. arenosa*, 2) trans-specific polymorphism in *A. arenosa* only (tetraploid SNP occurs in *A. arenosa* but not *A. lyrata* diploids and in one to all diploid congeners), 3) trans-specific polymorphism in *A. lyrata* only (tetraploid SNP occurs in *A. lyrata* but not *A. arenosa* diploids and in one to all diploid congeners), 4) trans-specific polymorphism in the congeners only, 5) *A. arenosa*-specific polymorphism, 6) *A. lyrata*-specific polymorphism, 7) tetraploid-private polymorphism. Here we differentiate between trans-specific and species-specific variability at the diploid level, reflecting ancestral allele sharing rather than recent introgression. It is crucial to note that any tetraploid SNP identified here ultimately represents trans-specific polymorphism since it is shared between tetraploids of both *A. arenosa* and *A. lyrata*.

As additional evidence to estimate variability in allelic sources of tetraploid haplotype blocks, we constructed genetic distance networks using Nei’s distance (Nei, 1972) in the ‘adegenet’ package in R (Jombart, 2008). We visualized these networks using SplitsTree (Huson & Bryant, 2001).

## Results and discussion

### Hybridization among tetraploid lineages of A. lyrata and A. arenosa

To reconstruct allelic sources of candidate WGD-adaptation haplotype blocks, we compiled a collection of genomes from 983 individuals: 818 individuals from 46 diploid and 61 tetraploid populations of *A. lyrata* and *A. arenosa*, and 165 individuals from four congeners. Our new sampling focused on Central European *A. lyrata* to cover all regions possibly harbouring the tetraploid cytotype (Ansell et al., 2010; Marburger et al., 2019; Schmickl & Koch, 2011; Seear et al., 2020) and a representative sampling of all European diploid outcrossing *Arabidopsis* species.

We observed that *A. lyrata* autotetraploids (‘tetraploids’ hereafter) occur at geographically distinct locations throughout Central Europe (Fig. 2, Dataset 1). To obtain comparable and representative samples of *A. lyrata* and *A. arenosa* populations for diploid-tetraploid selection scans, we first analyzed 154 individuals from 17 proximal diploid and tetraploid populations (Fig. 2). We assessed population structure and potential admixture (Fig. 2) using bootstrapped allele covariance trees (TreeMix; Fig. 2B, C), neighbor-joining networks, and Bayesian clustering (FastStructure; Fig. S1). In *A. arenosa*, we identified a single Central European tetraploid lineage, consistent with previous studies (Arnold et al., 2015; Monnahan et al., 2019). Notably, *A. lyrata* tetraploids consisted of three differentiated lineages in Austria, Czechia, and Germany. This suggests either independent formation and establishment of tetraploid populations in each region, or a single origin of autotetraploid *A. lyrata* followed by allopatry and, potentially, local introgression from parapatric diploid *A. lyrata* lineages. Both species and cytotypes maintained high genetic diversity (*A. arenosa*: mean nucleotide diversity over four-fold degenerate sites (4d-π) = 0.026/0.024 for diploids/tetraploids; *A. lyrata*: mean 4d-π = 0.012/0.016 for diploids/tetraploids, respectively; Table S1). Although Central European *A. lyrata* has a more scattered distribution and lower nucleotide diversity than *A. arenosa*, it showed neutral Tajima’s D (mean Tajima’s D in *A. arenosa* = 0.01/0.20 for diploids/tetraploids; mean Tajima’s D in *A. lyrata* = 0.27/0.23 for diploids/tetraploids; Table S1). Altogether, our analyses identified three tetraploid lineages in Central European *A. lyrata*, in addition to the single well-described tetraploid lineage in *A. arenosa*.

**Fig. 2:**
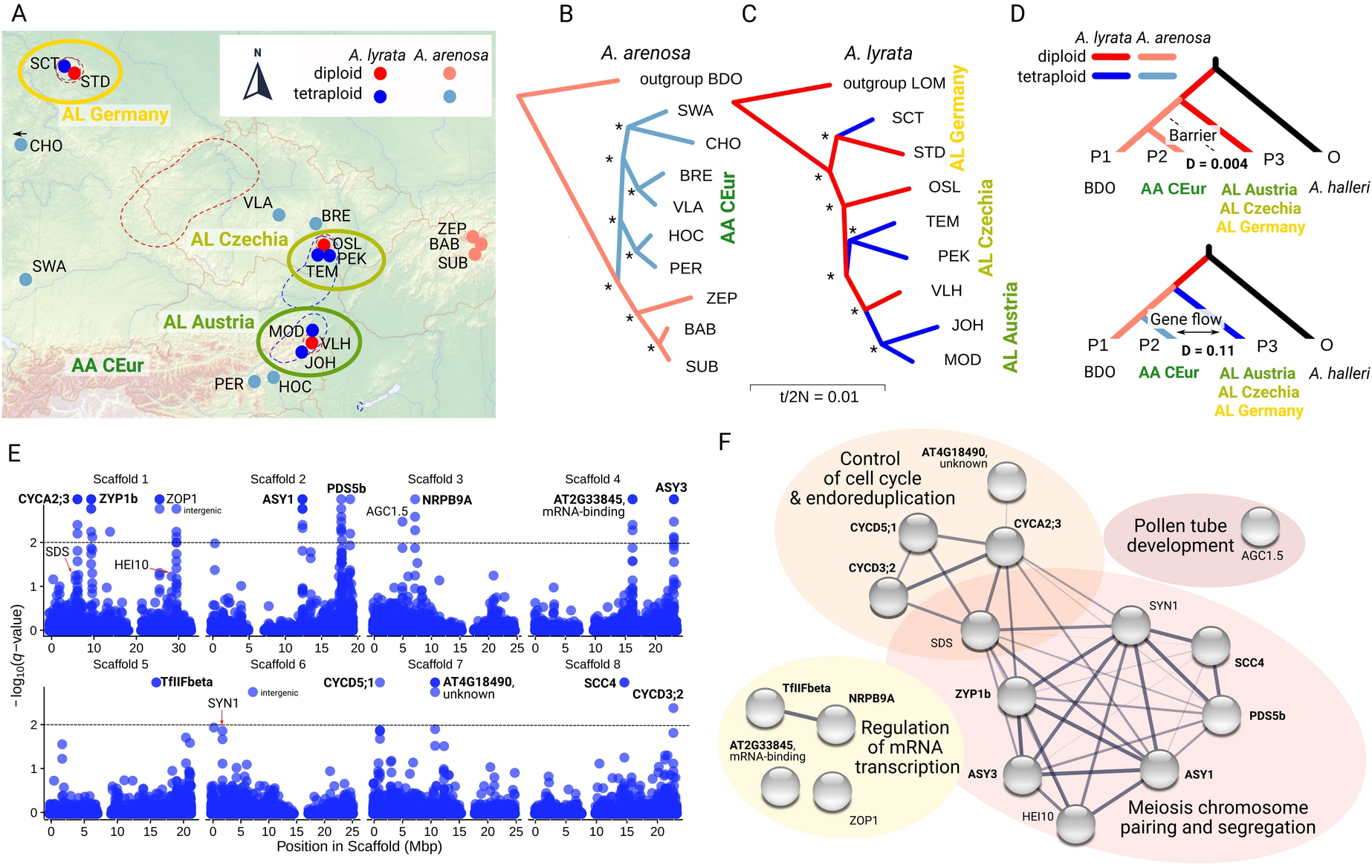
Evolutionary relationships and genomic signatures of selection in tetraploid populations of A. lyrata and A. arenosa. A: Locations of our focal 17 populations (AL Austria, AL Czechia, and AL Germany) and A. arenosa (AA CEur) in Central Europe. Red and blue dashed lines show the ranges of diploid and tetraploid A. lyrata, respectively; tetraploids of A. arenosa occur throughout the entire area. B, C: Phylogenetic relationships among the focal populations of A. arenosa (B) and A. lyrata (C) inferred by TreeMix analysis. Asterisks show bootstrap support = 100%. D: Introgression among tetraploid but not diploid populations of both Arabidopsis species. ABBA-BABA analysis demonstrating excess allele sharing between tetraploids (bottom tree), but not diploids (top tree) of A. arenosa and each of the three A. lyrata tetraploid lineages. P1 is BDO, the earliest diverging and spatially isolated diploid population of A. arenosa (Kolář, Fuxová, et al., 2016), and outgroup is A. halleri from Austria. E: A set of 14 unlinked genes showing significant evidence (p < 0.01) of positive selection (positively selected genes, PSGs) in the four tetraploid lineages identified using PicMin. Additional three genes, SDS, HEI10, and SYN1, were identified using a screen for candidate SNPs. F: Functional characterization of the 17 PSGs by STRING analysis. The network shows predicted protein-protein interactions among the PSGs. The width of each line corresponds to the confidence of the interaction prediction. PSGs were annotated into four processes, each represented by a bubble of different colour. The 12 PSGs with names written in bold had enough candidate SNPs for the reconstruction of tetraploid haplotype blocks (see the main text).

Next, we examined introgression between the two species using D statistics (ABBA-BABA, (Martin et al., 2013)). While insignificant among diploids, we identified evidence of introgression (D between 0.106 to 0.108; Fig. 2D) between tetraploids of *A. arenosa* and each of the three tetraploid *A. lyrata* lineages. These results are in line with previous experimental studies: while diploids of both species exhibit strong postzygotic barriers, polyploidy-mediated hybrid seed rescue enables hybridization between tetraploid *A. arenosa* and *A. lyrata* (Lafon-Placette et al., 2017).

In summary, we identified four tetraploid lineages in Central European *Arabidopsis*, consistent with two to four independent WGD events (one in each species and possibly up to two additional WGD events in *A. lyrata*). These WGD events reduced between-species hybridization barriers (Lafon-Placette et al., 2017), leading to interspecific gene flow between tetraploids (Marburger et al., 2019; Schmickl & Koch, 2011). This makes hybridization among also our novel polyploid lineages a plausible source of putatively beneficial alleles in this tetraploid system.

### Genomic signatures of repeated differentiation associated with WGD

To infer candidate genes associated with adaptation to WGD, we compared each of the four tetraploid *A. lyrata*/*A. arenosa* lineages to their geographically most proximal diploids (Fig. 2B, C). We identified candidate genes using PicMin, a novel method that identifies repeated signatures of selection (Booker, 2023). We identified 54 significantly differentiated windows (PicMin FDR-corrected *q*-value < 0.01; Dataset S2), overlapping 14 unlinked candidate genes (Fig. 2E, Table S2). PicMin measures across windows; to capture narrower peaks of differentiation, we also conducted a search for genes featuring highly diploid/tetraploid differentiated SNPs present in multiple lineages (see Methods). Such a ‘candidate SNP’ approach can identify genes undergoing positive selection in tetraploids on a subset of SNPs while the evolution of the rest of the gene sequence is constrained. Such analysis, with 1% outlier Fst cutoff, identified all 14 PicMin candidate genes (Table S2) and three additional candidates (HEI10, SYN1, and SDS), each with multiple outlier SNPs differentiated across all four tetraploid lineages (Dataset S3). In total, we recognized 17 candidate positively selected genes (PSGs) showing robust, repeated selection signals in tetraploid lineages. Notably, these loci do not cluster in regions characterized by extreme recombination rates, indicating no bias associated with the recombination landscape (Fig. S2 (Burri, 2017)).

To understand in which molecular processes the 17 WGD-associated PSGs are involved, we predicted protein-protein interactions among them using STRING analysis (Szklarczyk et al., 2015). We retrieved two interconnected clusters: regulation of cell cycle and endoreduplication by cyclins, and chromosome pairing and segregation during meiosis (p < 0.001, Fisher’s exact test; Fig. 2F). To investigate the phenotypic impact of PSGs associated with the cell cycle trait endoreduplication on tetraploids, we employed flow cytometry. Our findings revealed a substantial decrease in endoreduplication within established tetraploids compared to newly synthesized tetraploids, suggesting compensatory evolution leading to reduced DNA content through decreased endoreduplication (Supplementary Text 2, Fig. S4, S5). This is consistent with results of cytological studies reporting stable meiotic chromosome segregation in established tetraploids of *A. arenosa* and *A. lyrata*, compared to newly synthesized tetraploids of *A. arenosa*, which suggests a compensatory shift in the meiotic stability phenotype (Marburger et al., 2019; Morgan et al., 2022, 2020; Seear et al., 2020; Yant et al., 2013). Thus, the interacting meiosis and cell cycle regulation PSGs likely mediate beneficial phenotypic shifts resulting in the establishment of tetraploid lineages.

Despite the high overall functional connectivity, six of the 17 PSGs were not connected in this interaction network. Interestingly, four of these six genes were found to be related to mRNA transcription via RNA polymerase II (Fig. 2F) (TAIR database (Berardini et al., 2015), last accessed 02/03/2023). The fifth tetraploid PSG (*AT4G18490*) encodes an unknown protein which is strongly expressed in young *A. thaliana* flower buds and involved in cell division (Berardini et al., 2015). The sixth gene, AGC1.5, was not connected to any of the above-mentioned processes, but regulates pollen tube growth (Zhang et al., 2009). We provide further functional interpretations in Supplementary Text 1 and Fig. S3.

To summarise, the 17 tetraploid PSGs mediate processes of homologous chromosome pairing during prophase I of meiosis, cell cycle timing and regulation of endoreduplication via different classes of cyclins, and mRNA transcription via RNA polymerase II.

### Novel tetraploid haplotype blocks are assembled from diverse allelic sources

Both theoretical and empirical evidence indicates that polyploids may have increased access to allelic variation than diploids (Arnold et al., 2016; Monnahan et al., 2019; Ronfort, 1999; Selmecki et al., 2015). Here we inquired if these positively selected regions accreate as novel mosaics from different allelic sources (Fig. 1B). To do this, we first deconstructed haplotype blocks in our PSGs and identified the specific allelic sources for every candidate tetraploid SNP. Removing five PSGs with insufficient variation (less than five candidate SNPs), we analysed 12 genes and 232 candidate SNPs (Dataset S4). For each gene, we reconstructed diploid- and tetraploid-characteristic candidate haplotype blocks using these candidate SNPs as markers (following (Bohutínská, Handrick, et al., 2021)). These haplotype blocks corresponded to real haplotypes observed in PacBio HiFi reads from five diploid and five tetraploid individuals of *A. arenosa* (71% correspondence across 132 checked reads, 95% correspondence if heterozygous sites were excluded; Dataset S4-S6).

We identified a single major tetraploid-characteristic haplotype block for each of the 12 candidate genes that was shared across both *A. lyrata* and *A. arenosa* tetraploids (Fig. 3B; present in 92 and 98% of populations, respectively; mean frequency = 0.62 across 61 tetraploid populations; Table S4). Based on estimates of the age of the tetraploids, which range from approximately 20,000 to 230,000 generations (Arnold et al., 2015; Marburger et al., 2019), we speculate that the major tetraploid haplotype block likely rapidly spread across Europe. We further identified two major diploid haplotype blocks for each gene, one *A. lyrata*-characteristic and one *A. arenosa*-characteristic (Fig. 3B). Other haplotype blocks were of minor frequency (Fig. 3B). Importantly, none of the tetraploid haplotype blocks associated with the 12 PSGs were found in any diploid individuals, even after a sampling that included the potential source lineages for the tetraploids. This indicates that they likely assembled upon the establishment of tetraploids rather than pre-existing as standing variation in diploids.

**Fig. 3:**
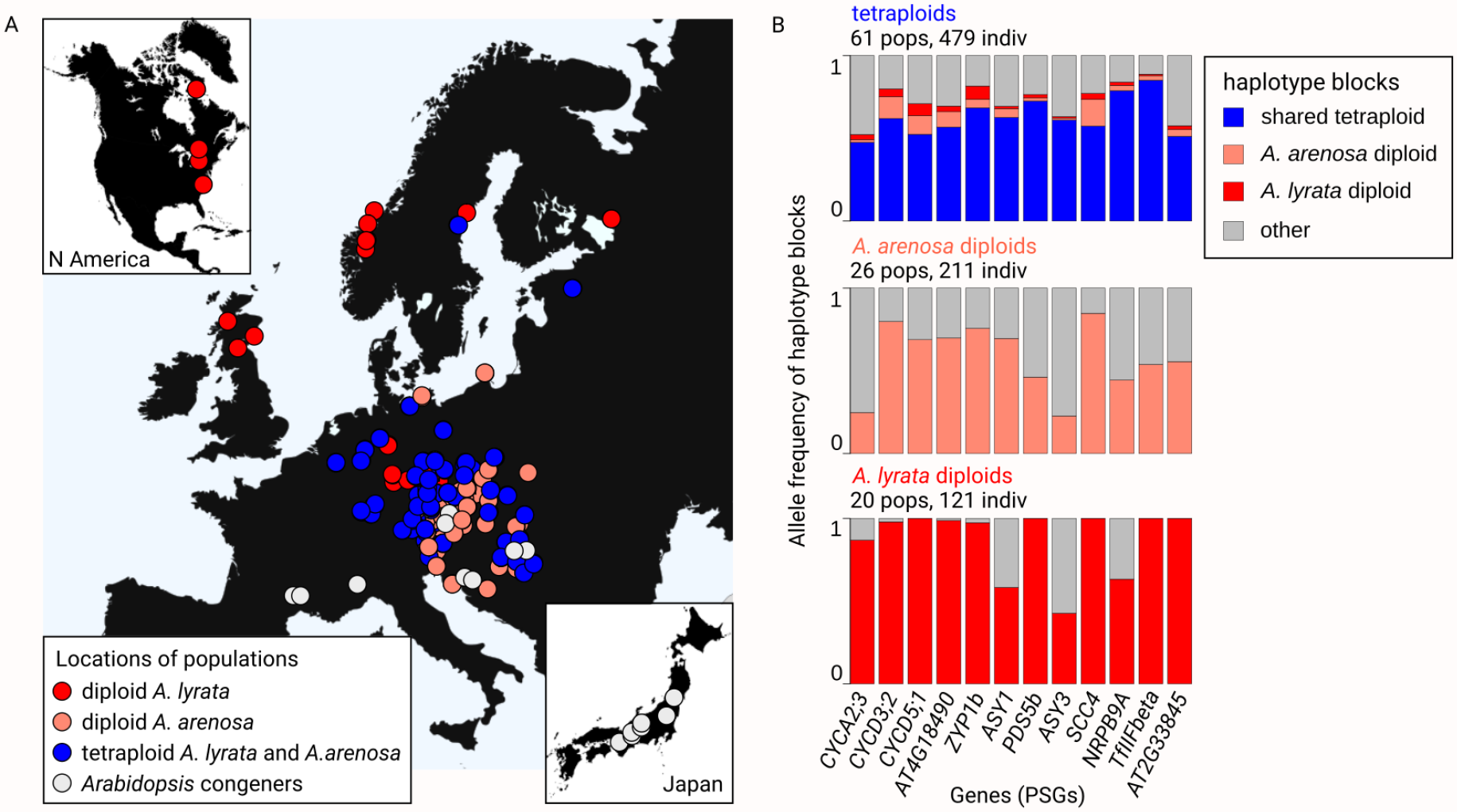
Distribution of candidate haplotype blocks. A: Populations in this analysis (504 diploids and 479 tetraploids). Arabidopsis congeners are A. halleri, A. croatica, A. cebennensis, and A. pedemontana. B: Total absence of tetraploid haplotype blocks in diploids. Frequency of shared tetraploid (blue), A. arenosa diploid (light red), and A. lyrata diploid (red) haplotypes in each of the 12 top candidate genes. All other haplotypes present (including possible recombinants of the above) are shown in grey.

Despite the absence of each of the 12 complete tetraploid haplotype blocks in diploids, certain tetraploid SNPs within these blocks were detected across diploid populations, present in one or even across multiple species (Fig. 4). We therefore investigated the possibility that the tetraploid haplotype blocks might be assembled from multiple sources. To do this, we inferred the most likely source for each candidate SNP marking these tetraploid haplotype blocks. Considering the extent of trans-specific polymorphism in *Arabidopsis* (Novikova et al., 2016), we worked with the full dataset of *A. arenosa, A. lyrata*, and all other diploid *Arabidopsis* outcrossers (*A. halleri, A. croatica, A. cebennensis*, and *A. pedemontana*) (Dataset S1). Using the phylogenetic relationships among these species (Novikova et al., 2018) and the SNP presence/absence data from all 504 diploid individuals, we identified the most probable source for each of the 232 SNPs defining haplotype blocks in the 12 PSGs (Fig. 4, Dataset S7). Surprisingly, 65.5% of the SNPs forming tetraploid haplotype blocks were most parsimoniously inferred as trans-specific polymorphism, segregating in diploids of multiple *Arabidopsis* species (Fig. 4A, scenarios 1-4). Note that this number is likely an underestimate as some alleles may have remained unsampled or have gone extinct in any diploid lineage. This supports the growing recognition of the role of trans-specific standing polymorphism in adaptation (Guggisberg et al., 2018; Marques et al., 2019)). Only 6.9% of the candidate SNPs were inferred as arising from standing variation from a single diploid progenitor (*A. arenosa* or *A. lyrata*; Fig. 4A, scenarios 5, 6), and the remaining 27.6% of SNPs possibly accumulated de novo in tetraploids or remained unsampled in our dataset (absent in any of the 504 diploid individuals; Fig. 4A, scenario 7). We further quantified that 35.8% of the candidate SNPs marking the tetraploid haplotype blocks were likely contributed from diploid *A. arenosa* (present in this species and possibly in diploids of other species, but not *A. lyrata*; Fig. 4A, scenarios 2, 5), while 3.4% likely came from diploid *A. lyrata* (Fig. 4A, scenarios 3, 6). Finally, in the process of tetraploid haplotype block formation through these identified source scenarios, 71.1% of the candidate SNPs were indicative of likely introgression across all four tetraploid lineages. This was inferred from their presence in tetraploids and in at most one of the diploid progenitors (either diploid *A. arenosa* or *A. lyrata;* note that this scenario is nested within others; Fig. 4A, scenarios 2-7), aligning with the observed signatures of hybridization among tetraploids of both species (Marburger et al., 2019).

**Fig. 4:**
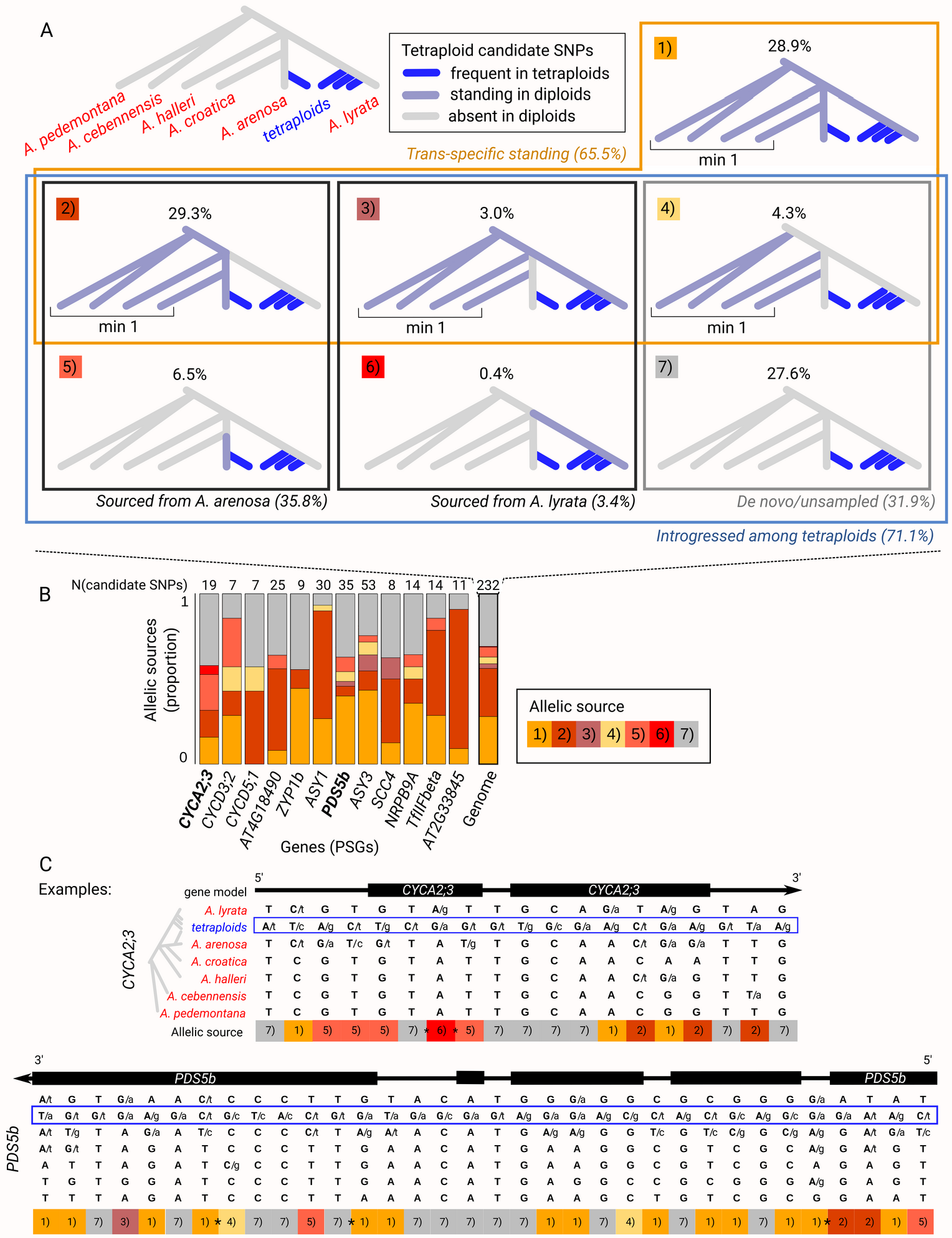
Mosaic of allelic sources of tetraploid haplotype blocks. A: The 232 candidate SNPs marking the 12 tetraploid haplotype blocks were categorized into one of the seven source scenarios based on their allele distribution in the 983 samples. The most parsimonious origin of each pattern is provided in italics. Four (outlined by orange frame) involve trans-specific standing variation shared among diploids of at least two species. Six (outlined by blue frame) require introgression between tetraploid lineages. Boxes ‘min 1’ show that the tetraploid standing variation is present in at least one congener. Phylogenetic relationships according to (Novikova et al., 2018). B: Variable source scenarios for each of the 12 tetraploid positively selected genes (PSGs). Barplots show the proportion of candidate SNPs representing each of the seven source scenarios, numbers above bars indicate the number of candidate SNPs per each PSG. Bold highlights two example PSGs shown at panel C. C: Illustration of diploid and tetraploid haplotype blocks for two PSGs. Bold capital letters represent haplotype-marking alleles, and two letters at a position indicate the presence of an alternative minor allele. Coding sequences highlighted as boxes (only regions overlapping with candidate SNPs are shown). The bottom line displays SNP assignment to its source scenario as defined in panel A. Possible recombination breakpoints required to construct the haplotype from various allelic sources are marked with asterisks. Top: example from the CYCA2;3 endoreduplication gene, displaying 19 candidate SNPs spanning 5371 bp. Bottom: example from the PDS5b meiosis gene, depicting candidate SNPs spanning 8904 bp. Description is consistent with the top panel.

The different allelic sources forming tetraploid haplotype blocks varied genome-wide (when analysing all 12 PSGs together; Fig. 4A), but also within individual tetraploid haplotypes (3-6 scenarios per haplotype, median = 4, out of 7; Fig. 4B, C, Fig. S6). This suggests that tetraploid haplotype blocks represent a combination of reuse of species-specific as well as trans-specific SNPs, with additional input of de novo mutations after WGD (Fig. 4B, C). Such composite ‘mosaic’ haplotype blocks were likely spread among tetraploid lineages by introgression. The rearrangement of diverse allelic sources into mosaic tetraploid haplotypes may be facilitated by high recombination rates reported for these *Arabidopsis* species (Dukić & Bomblies, 2022; Hämälä & Savolainen, 2019) and the observation of enhanced recombination rates in neo-tetraploid *A. arenosa* and *A. thaliana* (Morgan et al., 2020; Pecinka et al., 2011). The mosaic nature of allelic sources was further supported by the reticulation observed in the neighbor-joining networks of the 12 PSGs (Fig. S7). In these networks, tetraploids form a single lineage that connects through multiple splits to two or more diploid *Arabidopsis* lineages. We provide hypotheses about the spatio-temporal context of the origin of these mosaic haplotypes in Supplementary Text 2.

In summary, our analysis of variants forming tetraploid haplotype blocks indicates that polyploids acquire alleles from diverse source pools, combine them during post-WGD adaptation, and further redistribute this variation by gene flow among polyploid lineages. Our results also highlight the role of interspecific introgression in generating potentially beneficial diversity within polyploid species complexes, complementary to earlier studies (reviewed in (Leal et al., 2023, Marhold & Lihová, 2006; Schmickl & Yant, 2021)).

### Concluding remarks

Here we investigated how candidate WGD-adaptive haplotypes may form in autopolyploids. Leveraging our very deep sampling of all extant outcrossing *Arabidopsis* species in Europe, we identified a set of genes involved in cell cycle, meiosis, and transcription that consistently exhibit the strongest signals of selection. All tetraploids reused identical haplotype blocks across the diagnostic markers, but these blocks were entirely absent from diploids. Importantly, each haplotype block comprised SNPs of broadly varying ancestry, including trans-specific and species-specific polymorphisms from closer and more distant relatives, along with likely de novo mutations in the tetraploids. These ‘evolutionary mosaic’ haplotype blocks were then likely shared via introgression among tetraploids and spread across Europe.

Altogether, we demonstrated that extensive re-shuffling of trans-specific, species-specific, and novel variation occurred in response to the challenges of WGD. Our study demonstrates that polyploid lineages leverage their enhanced capacity to accumulate genetic variation from various sources to generate new, potentially advantageous haplotypes. Importantly, this process is not unique to polyploids, as diploids have also been reported to accumulate multiple adaptive changes into finely tuned haplotypes (Archambeault et al., 2020; McGregor et al., 2007; Roberts Kingman et al., 2021). Recent research in various plant and animal species has revealed substantial variation in the contributions of introgressed, standing, and de novo alleles to adaptation at the entire gene level (Bohutínská, Vlček, et al., 2021; Konečná et al., 2021; Moran et al., 2022; Wang et al., 2021). The diversity of allelic sources in individual haplotype blocks in tetraploid *Arabidopsis* implies that pathways to adaptation may be even more diverse when considering the assembly of individual variants into haplotypes.

## Acknowledgements

We greatly appreciate the constructive feedback from members of the Ecolgen team in Prague, Katie Peichel, Reto Burri, Alison Scott, Polina Novikova, Andrew MacColl, Tuomas Hämälä, John Brookfield, Emma Curran, Laura Dean, Sian Bray, and Ana da Silvia. We further appreciate inspiration provided by the ForBio Polyploid course in Drøbak, Norway. We thank Martin Čertner and Dorka Čertnerová for sharing their polyploid synthesis protocol, Bodo Schwarzberg (Untere Naturschutzbehörde Nordhausen) for sharing seeds of the STD population, and Bertram Preuschhof (Untere Naturschutzbehörde Göttingen) for a collecting permit for the SCT population.

## Data and code availability

Sequence data that support the findings of this study are deposited in the NCBI (https://www.ncbi.nlm.nih.gov/bioproject/) under BioProjects PRJNA284572, PRJNA309929, PRJNA357693, PRJNA357372, PRJNA459481, PRJNA493227, PRJEB34247 (ENA), PRJNA506705, PRJNA484107, PRJNA592307, PRJNA667586, PRJNA929698. See

Dataset S1 for individual codes.

ScanTools_ProtEvol pipeline: github.com/mbohutinska/ScanTools_ProtEvol ABBA-BABA pipeline: github.com/simonhmartin/tutorials/tree/master/ABBA_BABA_whole_genome)

PicMin: github.com/TBooker/PicMin

Allele frequencies of haplotype blocks: github.com/mbohutinska/repeatedWGD, section ‘Haplotype AF’

