## Supplementary for "Novel reassortments of variation underlie repeated adaptation to whole genome duplication in two outcrossing Arabidopsis species"

#### Supplementary Text 1

The 17 tetraploid positively selected genes (PSGs) identified in this study are involved in a range of processes, including homologous chromosome pairing during prophase I of meiosis, cell cycle timing and regulation of endoreduplication through the action of different types of cyclins, and mRNA transcription via RNA polymerase II.

In previous research, the role of meiosis genes in tetraploid adaptation has been identified and tested, with results suggesting that these functional changes contribute to the successful establishment of tetraploids (Yant et al. 2013; Morgan et al. 2020; Seear et al. 2020; Morgan et al. 2022). Cell cycle regulation has also been studied in relation to polyploidy. Cyclins D1 and D2 have been shown to promote tolerance to genome doubling in tumors (Potapova et al. 2016; Crockford et al. 2017), while cyclin A2;3 regulates the extent of endoreduplication in *A. thaliana* (Imai et al. 2006), and cyclin D5;1 determines the rate of endoreduplication (Sterken et al. 2012). These findings suggest that cyclins may play a role in the adaptation to whole genome duplication (WGD), potentially by regulating endoreduplication levels.

Interestingly, positive selection acts at different types of SNPs among these three processes. The selection of upstream regulatory variability was more prevalent among cyclins (45% of upstream regulatory SNPs in cyclins compared to 32% genome-wide, $p = 0.16$, Chi square test). In contrast, selection acting on missense SNPs in coding regions was more common in meiosis genes (40% of missense SNPs in meiosis genes compared to 28% genome-wide, $p = 0.001$, Chi square test). Finally, transcription genes showed positive selection on synonymous SNPs in coding regions (65% of synonymous SNPs in transcription genes compared to 27% genome-wide, $p < 0.001$, Chi square test) (Fig. S6). These results suggest that the evolution of cyclins may have a more regulatory, possibly dosage-dependent basis, while meiosis evolution may be driven more by structural protein changes. It remains to be determined if the repeated differentiation in synonymous positions of some of the transcription genes results in any functional changes.

To summarize, our research has provided evidence for the involvement of cyclins, meiosis genes, and transcription-regulating genes in the adaptation to WGD. Cyclins may play a role in regulating the level of endoreduplication, while meiosis genes may be involved in structural protein changes. The role of transcription-regulating genes in the adaptation to WGD has also been proposed (Hollister et al. 2012; Marburger et al. 2019), and the transcription cycle was shown to be linked with the cell cycle, through the interaction of RNA polymerase II with cyclin-dependent kinases (Bregman et al. 2000; Guo and Stiller 2004). It is possible that these genes interact to jointly re-time the cell cycle in response to WGD.

#### Supplementary Text 2

Some phenotypic shifts in polyploids have been associated with the processes identified in our protein interaction analysis (Fig. 2F). Specifically, cytological studies reported stable meiotic chromosome segregation in established tetraploids of *A. arenosa* and *A. lyrata* compared to neo-tetraploids of *A. arenosa*, which suggests a compensatory shift in the meiotic stability phenotype (Yant et al. 2013; Marburger et al. 2019; Morgan et al. 2020; Seear et al. 2020; Morgan et al. 2022). Here, we extended the inquiry of polyploidy-related trait shifts to cell cycle phenotypes, asking if established tetraploids reduce their level of endoreduplication, as reported in *A. thaliana* and alpine lineages of *A. arenosa* (Imai et al. 2006; Sterken et al. 2012; Wos et al. 2022). To do so, we grew plants from each of the four established tetraploid lineages, their closely related diploids, and synthetic (oryzalin-induced) neo-tetraploids. We found that the level of endoreduplication (the fraction of endoreduplicated nuclei) and the number of endoreduplication cycles were consistently higher in diploids compared to tetraploids (Fig. S4). Notably, established tetraploids showed reduced endoreduplication compared to neo-tetraploids. Thus, the decrease of endoreduplication observed in tetraploids is likely due to post-WGD adaptation, rather than a mere phenotypic consequence of WGD itself. We found this decrease of tetraploid endoreduplication in each of the four lineages (Fig. S4, and p-values herein), and also when comparing all diploids with all neo-tetraploids and established tetraploids (mean level of endoreduplication = 0.82/0.74/0.61 for diploids/neo-tetraploids/tetraploids, $p < 0.001$; median maximum number of endoreduplication cycles = 3/2.5/2 for diploids/neo-tetraploids/tetraploids, $p < 0.001$; Wilcoxon rank sum test). In diploids, we observed a nonsignificant difference in the number of endoreduplication cycles (median = 3 in all four diploid lineages, $p > 0.05$, Wilcoxon rank sum test) and only a subtle difference in the level of endoreduplication in one lineage (Fig. S5). This suggests that the regulation of endoreduplication in *Arabidopsis* is generally conserved, as long as the intrinsic cell conditions do not change, for example as a result of WGD.

In summary, these results point to an adaptive compensation for DNA content per nucleus: while nearly double in neo-tetraploids, it returns toward diploid levels in established tetraploids (Fig. S4). This suggests ‘compensation’ over ‘trait change’ scenarios, as have been suggested for other polyploidy-specific traits following WGD (Bomblies 2020). Importantly, some of our strongest PSGs are associated with endoreduplication level regulation in *A. thaliana* or genome doubled tumours (genes CYCA2;3 (Imai et al. 2006), CYCD5;1 (Sterken et al. 2012)). Thus, these PSGs are possibly responsible for the reduction of endoreduplication in tetraploid *A. arenosa* and *A. lyrata*.

*Methods:*

To investigate whether genetic changes in cyclin genes may lead to a reduction in the level of endoreduplication, we grew 10 plants from each of the four lineages, from a diploid and tetraploid population each per lineage (totaling 80 plants), and we cultivated nine oryzalin-induced neo-tetraploid plants of *A. arenosa*, generated from the Western Carpathian diploid lineage, which was shown to be the progenitor of tetraploids (Arnold et al. 2015). We determined the endoreduplication profile of these plants using flow cytometric analysis on nuclei isolated from fully grown leaves (11th leaf in the rosette, (Wos et al. 2022)).

Specifically, we collected seeds from both diploid and tetraploid populations of *A. lyrata* and *A. arenosa* corresponding to their four tetraploid lineages and stored them at 4°C for vernalization. We germinated the seeds on moist filter paper in petri dishes. For induction of synthetic polyploids, we applied a drop of oryzalin solution (5% aqueous DMSO) to the shoot apical meristems of young seedlings at the stage of fully developed cotyledons. After 48 hours, we rinsed the seedlings with distilled water and transplanted them to soil, cultivating them under controlled conditions (20°C/16h day, 15°C/8h night).

To estimate the success rate of the oryzalin treatment, the ploidy level of each plant was tested by flow cytometry following (Doležel et al. 2007). Specifically, 0.5 $cm^2$ of leaf tissue (without middle vein) was chopped, using a razor blade, in Otto I buffer simultaneously with the same amount of *Carex acutiformis* as internal standard. Samples were then dyed with a staining solution consisting of Otto II buffer, DAPI (4',6-diamidino-2-phenylindole) and β-mercaptoethanol, and analysed on a CytoFLEX S flow cytometer (Beckman Coulter, Inc) equipped with a UV laser (375 nm, 60 mW) using the Beckman Coulter CytExpert Acquisition and Analysis Software v2.5. Only the plants with no or significantly low percentage (<5% of all events) of diploid nuclei (also showing no distinguishable population of nuclei) were considered good candidates for synthetic polyploids, and nine of them were chosen for further analysis.

To assess the level of endopolyploidy, the amount of DNA was estimated using a Partec CyFlow SL flow cytometer equipped with a 532 nm solid-state laser (Cobolt Samba 150 mW). Samples were processed using the same simplified two-step method with Otto buffers and stained with propidium iodide. Samples were analysed without internal standard. Resulting histograms/dot plots were evaluated using Partec FloMax v2.4d.

We calculated the level of endoreduplication as the number of endoreduplicated nuclei compared to the number of all nuclei in the analysis. We also quantified the maximum number of endoreduplication cycles that nuclei of each leaf underwent, considering only peaks supported by at least five nuclei of a given genome size. Finally, we calculated the endoreduplication intensity, measured as the weighted number of endoreduplication cycles per nucleus [$EI = (0 \times \%2C) + (1 \times \%4C) + (2 \times \%8C) + (3 \times \%16C) + (4 \times \%32C)$] (Sterken et al. 2012). We tested for the difference between diploid, neo-tetraploid, and tetraploid endoreduplication levels and numbers using the Wilcoxon test in the R package ‘stats’.

#### Supplementary Text 3

The finding of mosaic sources of tetraploid haplotypes raises questions on the spatio-temporal context of the origin of these haplotype blocks, in particular about historical area contacts of both species, hybridisation, and the timing of tetraploid establishment. Based on the available knowledge on the evolutionary history of both species and our results, we propose the following hypothesis. After their origin and primary establishment, tetraploid *A. lyrata* lineages may have exhibited rather minor signs of adaptation in the candidate genes that are in focus of our study, sweeping its adaptive diploid variability (3.4% of the total adaptive variation; Fig. 4A, scenario 3 and 6) plus an unknown portion of de novo mutations (Fig. 4A, scenario 7). In contrast to *A. lyrata*, tetraploid *A. arenosa* was found to bear a significant amount of likely adaptive variability inherited from its diploid progenitor (35.8% of the total variability is of *A. arenosa* diploid origin; Fig. 4A, scenario 2 and 5). In addition, tetraploid *A. arenosa* also adapted from an unknown portion of de novo mutations (Fig. 4A, scenario 7), altogether constituting a majority of the current adaptive haplotype blocks. *Arabidopsis arenosa* likely polyploidised in the Western Carpathians (Arnold et al. 2015; Monnahan et al. 2019), which is around 350 km apart from the current distribution range of *A. lyrata* (the spatially closest are the Czech and Austrian populations). This, and the fact that the primarily (sub)arctic species *A. lyrata* may have been more widespread under colder climate, could have led to peripatric hybridisation of both species during the large Pleistocene vegetation turmoil at the glacial/interglacial boundaries in the area (Abraham et al. 2016). Tetraploid-adaptive variants of tetraploid *A. lyrata* and *A. arenosa* may then have been introgressed in both directions (although introgression from *A. arenosa* to *A. lyrata* was found to be much more frequent) and recombined into the final tetraploid haplotype blocks, resulting in the currently observed mosaic haplotypes comprising SNPs of both *A. lyrata* and *A. arenosa* ancestry. These final tetraploid haplotype blocks likely offered a higher fitness advantage than the species-specific haplotypes, because they became pervasive in all tetraploid lineages of both species (Fig. 3B) and spread across the entire natural distribution range of autotetraploid *Arabidopsis* in Europe (Fig. 3A).

To formally test among these scenarios, we propose detailed population genomic sampling across all four tetraploid lineages and their contact zones, coupled with hierarchical demographic inference, to identify the most likely evolutionary history of this intriguing reticulate adaptation.

Further, it is a pending question why there is almost an order higher contribution of candidate SNPs of *A. arenosa* than *A. lyrata* ancestry (Fig. 4A, scenario 2 and 5 vs. 3 and 6). It may partly be because diploid *A. arenosa* is more genetically variable then *A. lyrata* (pi = 0.026/0.012 for *A. arenosa*/*A. lyrata*; Table S2, (Marburger et al. 2019)). Alternatively, *A. lyrata* may exhibit more trade-offs between a diploid and tetraploid fitness optimum, although we miss any evidence supporting this idea. Finally, there might have been a higher initial proportion of SNPs of *A. lyrata* ancestry, but adaptive variation inherited from diploid *A. arenosa* or originated de novo might have presented a higher selective advantage. Also, some SNPs of *A. lyrata* origin might have become extinct in diploid *A. lyrata* after WGD, and thus being categorized as de novo or unsampled origin (this category of candidate SNPs is classified into scenario 4 and 7, Fig. 4A). To answer if standing variation inherited from diploid *A. lyrata* brought a substantial improvement in the overall adaptation of tetraploids, it would help to understand the functional impact of *A. lyrata*-sourced variation. Although this goes beyond the scope of this study, we found that three of the seven likely *A. lyrata*-sourced mutations are nonsynonymous (Asp168Gly in CYCA2;3, Ser538Pro and Met482Thr in ASY3) while four are synonymous (in PDS5b, SCC4, and ASY3), suggesting functional impact of some, but not all of the variants.

#### Supplementary Figures

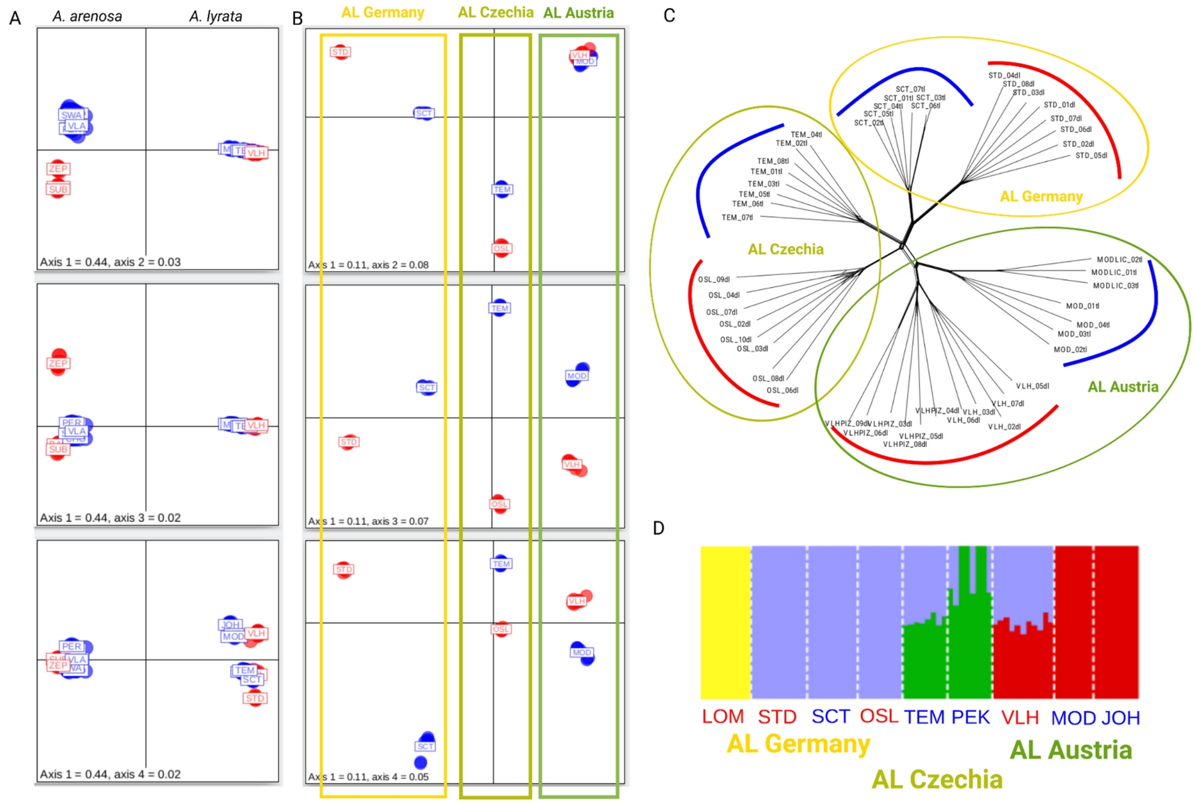


Fig. S1: Population structure of *A. arenosa* (A) and *A. lyrata* (A-D). A: PCA shows separation between *A. arenosa* and *A. lyrata* over 50,000 of scaffold_1 SNPs. (B-D): PCA, Nei's distance-based neighbour joining tree, and FastStructure plot all support three lineages of tetraploid *A. lyrata*, named AL Germany, AL Czechia, and AL Austria. For these analyses we used 1,094,553 genome-wide four-fold degenerate SNPs. Red: diploid populations, blue: tetraploid populations.

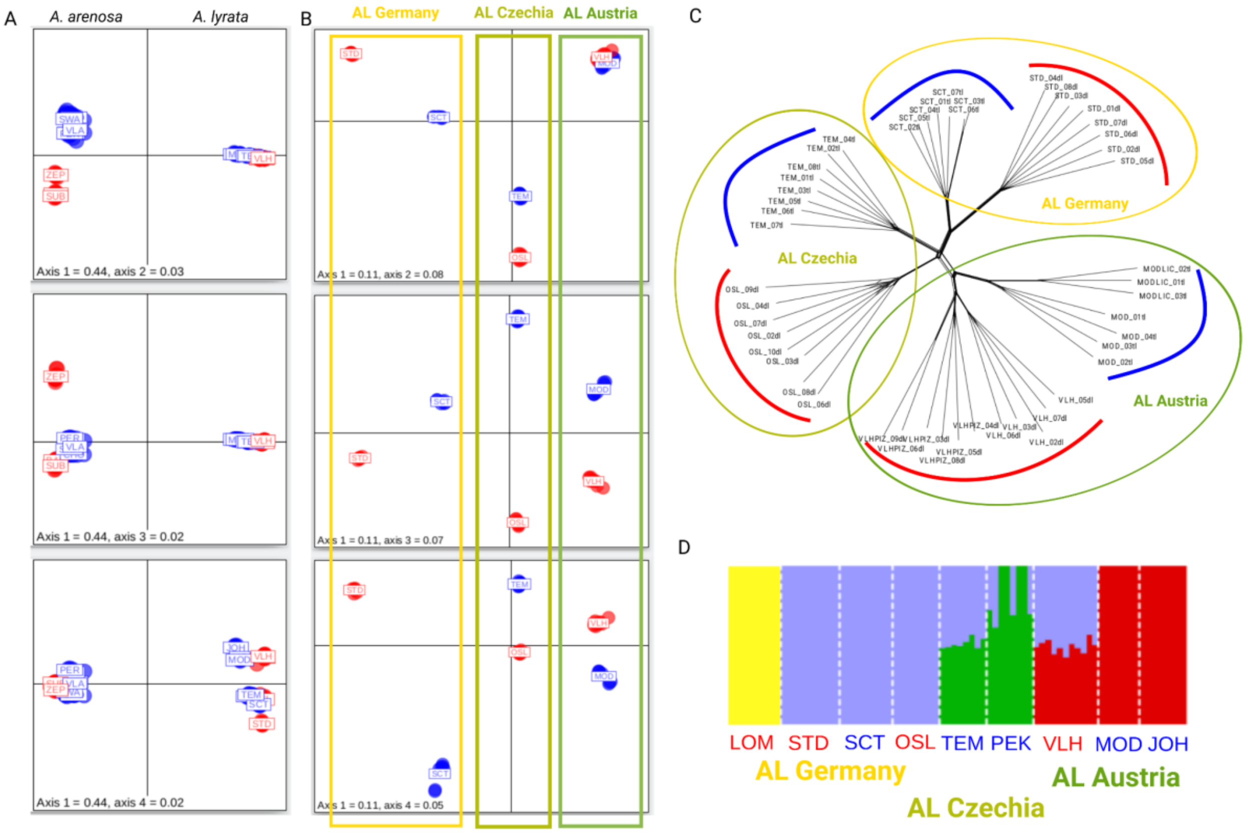

A
A. arenosa
A. lyrata
Axis 1 = 0.44, axis 2 = 0.03
Axis 1 = 0.44, axis 3 = 0.02
Axis 1 = 0.44, axis 4 = 0.02
B
AL Germany
AL Czechia
AL Austria
Axis 1 = 0.11, axis 2 = 0.08
Axis 1 = 0.11, axis 3 = 0.07
Axis 1 = 0.11, axis 4 = 0.05
C
AL Germany
AL Czechia
AL Austria
D
LOM
STD
SCT
OSL
TEM
PEK
VLH
MOD
JOH
AL Germany
AL Czechia
AL Austria

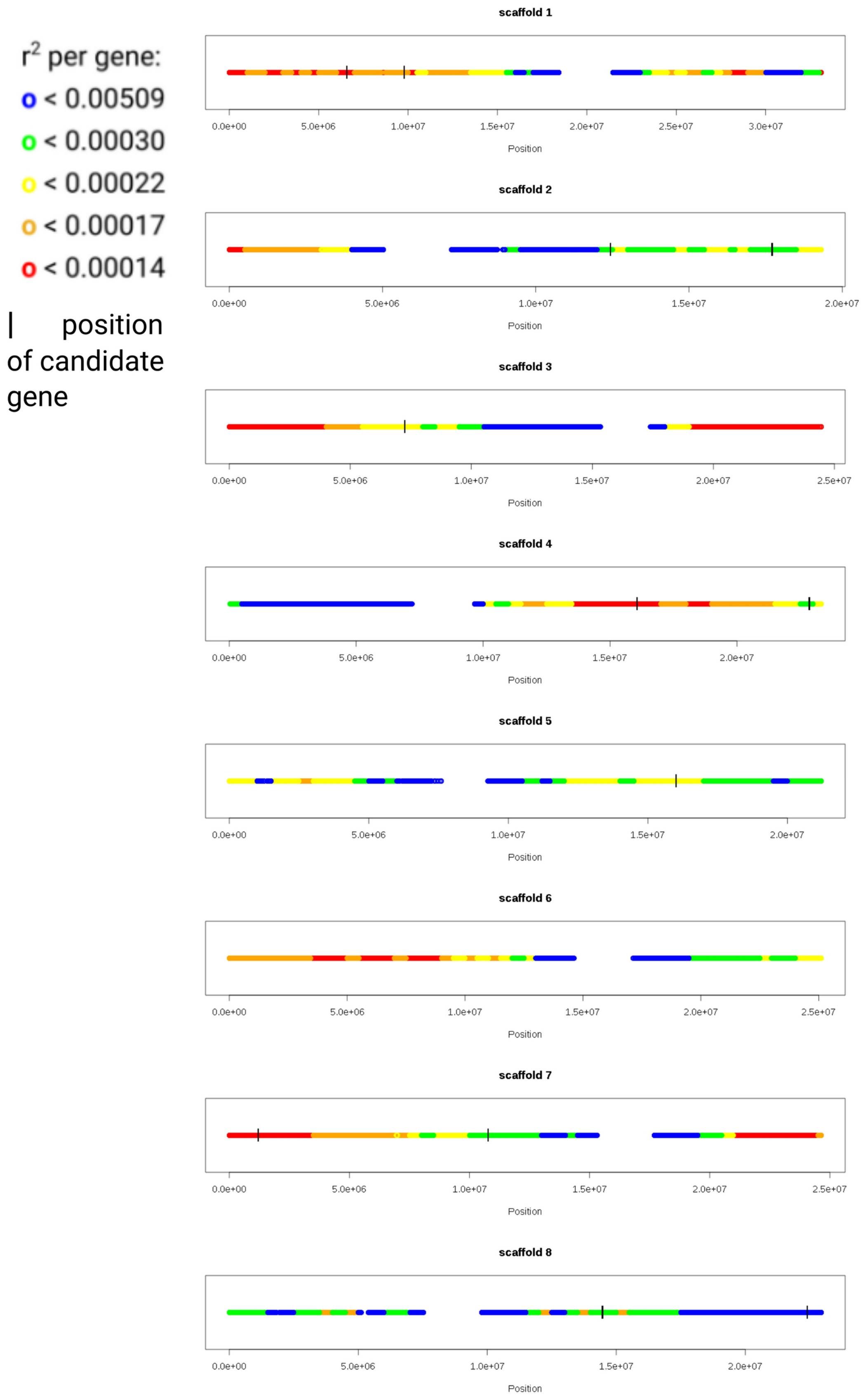


Fig. S2: Location of candidate genes (black vertical lines) on *A. lyrata* reference chromosomes colored by bins of distinct recombination rate per gene. The figure indicates that gene candidates do not cluster in regions with extreme values of recombination rate per gene (blue), as estimated based on the available *A. lyrata* recombination map (Hämälä and Savolainen 2019).

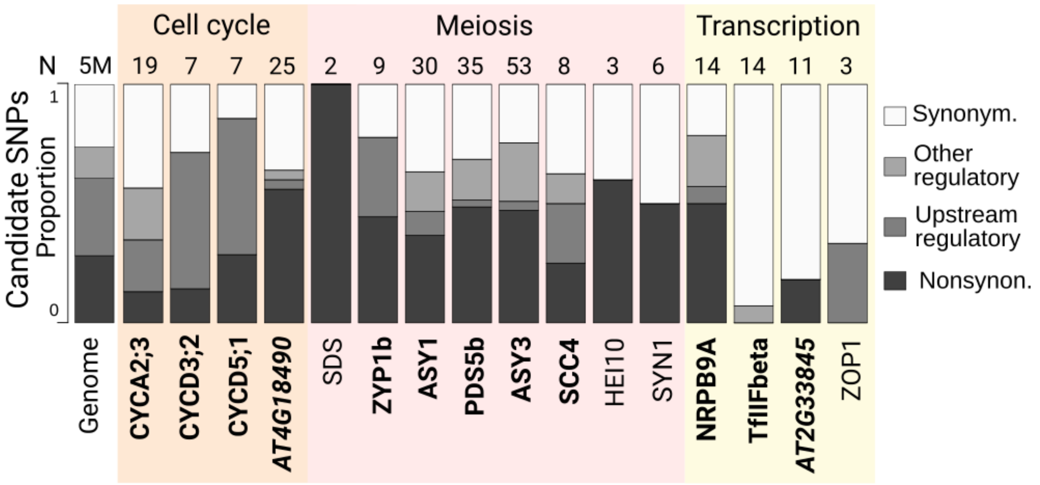


Fig. S3: Candidate PSGs involved in processes of cell cycle regulation, meiosis and transcriptional regulation vary in their proportions of differentiated cis-regulatory, nonsynonymous, and synonymous SNPs. The number above each bar corresponds to the number of differentiated SNPs within each gene. ~5M of genic and cis-regulatory SNPs from whole genome sequencing data were taken as controls.

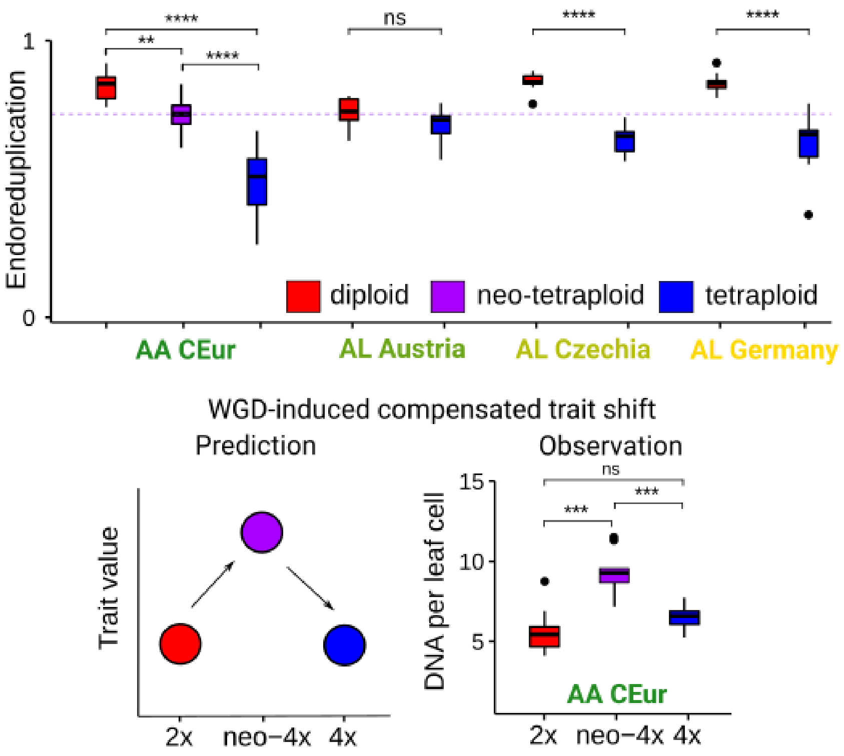


Fig. S4: Phenotypic shift associated with the establishment of tetraploids of *A. arenosa* and *A. lyrata* in the form of a decrease in the level of DNA endoreduplication. Top: Proportion of endoreduplicated nuclei in leaves of 10 individuals per each lineage and ploidy. **: $p < 0.01$, ****: $p < 0.0001$, ns: nonsignificant, Wilcoxon rank sum test. The horizontal violet line shows the level of endoreduplication under a scenario of no post-WGD adaptation, estimated from values of synthetic neo-tetraploids of *A. arenosa*. Bottom: Prediction for the compensatory evolution of polyploid traits from (Bomblies 2020) and observed average DNA content per leaf cell (calculated as a mean number of homologous chromosomes per nucleus) for diploid, neo-tetraploid, and tetraploid *A. arenosa*.

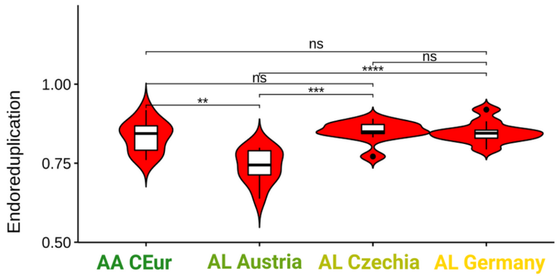


Fig. S5: Comparison of leaf endoreduplication among diploids. Upper boxplots show the endoreduplication level, calculated as the number of endoreduplicated nuclei divided by all nuclei in the analysis, bottom plots show the maximum number of endoreduplication cycles in the leaf per each diploid lineage. ****: p < 0.0001, ***: p < 0.001, ns: nonsignificant, Wilcoxon rank sum test. Each boxplot is represented by 10 individuals.

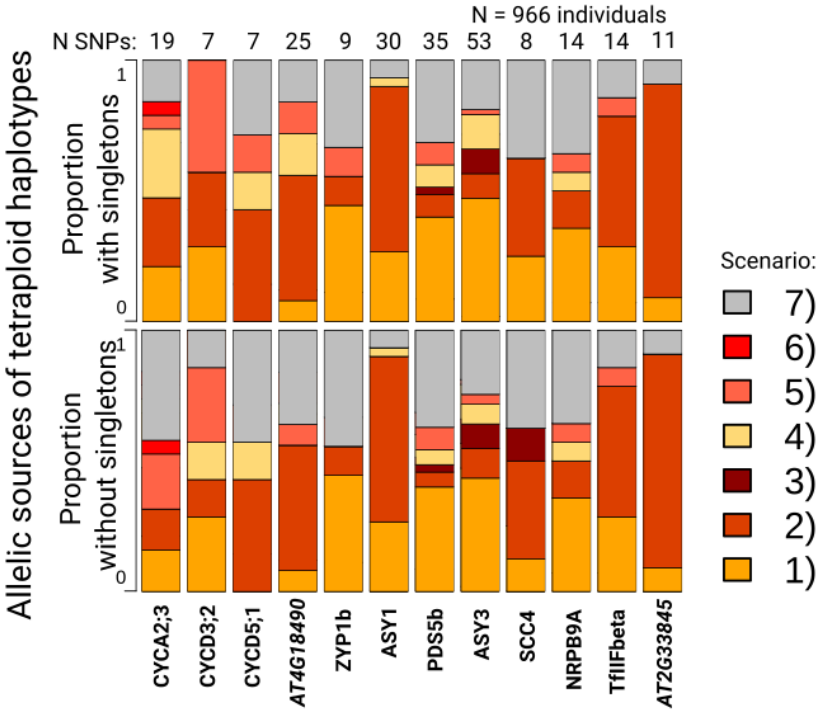


Fig. S6. Variable allelic sources of tetraploid haplotypes for each of the 12 tetraploid positively selected genes (PSGs). Barplots show the proportion of candidate SNPs representing each of the seven source scenarios (see Fig. 5 for graphical visualisation of scenarios). Upper plot shows the source when including singletons, bottom plot after filtering singletons out.

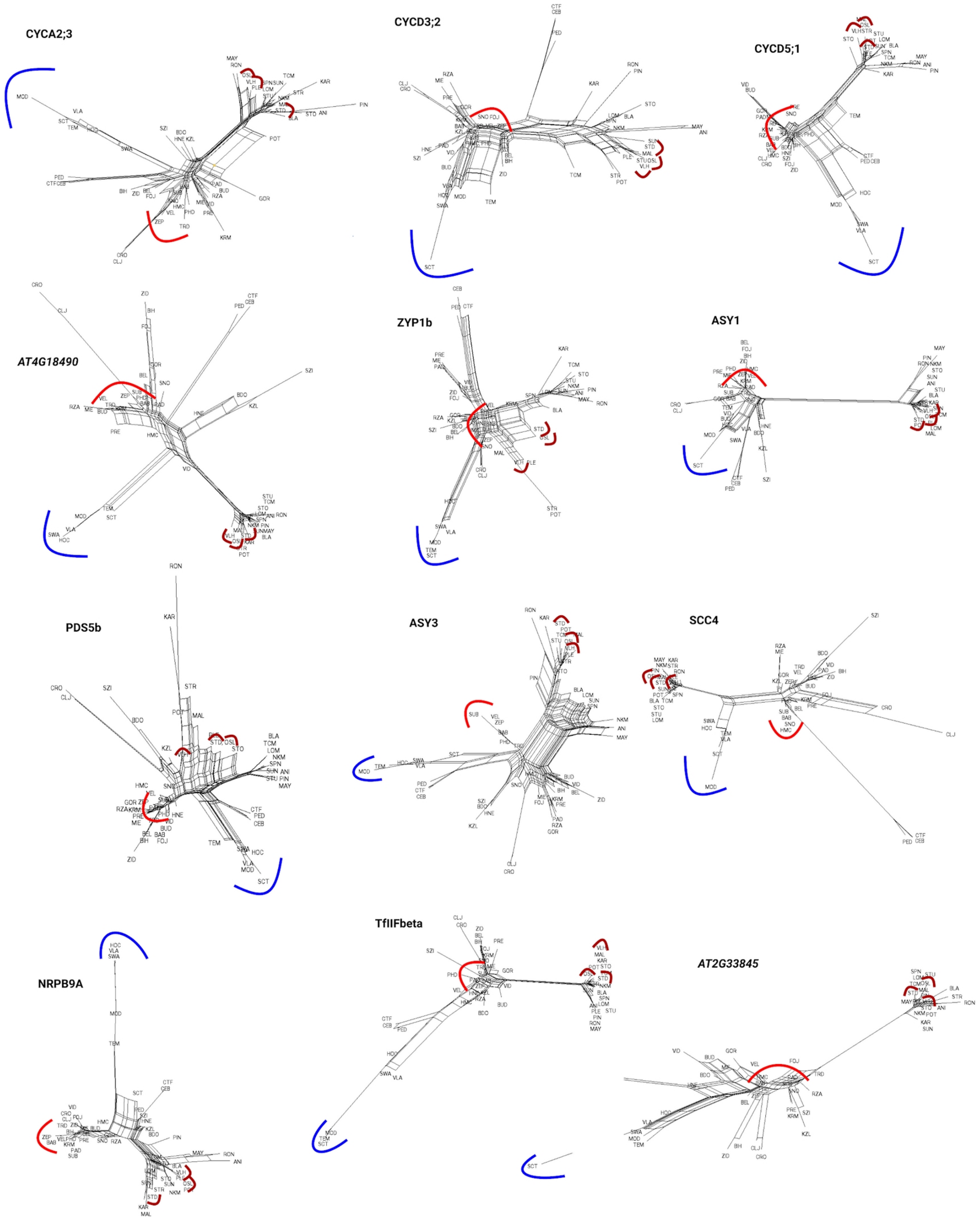


Fig. S7: Nei's distance-based neighbour-joining networks of tetraploid positively selected genes (PSGs) across diploid *Arabidopsis* (*A. arenosa*, *A. lyrata*, *A. croatica*, *A. cebennensis*, and *A. pedemontana*) and tetraploid *Arabidopsis* (*A. arenosa*, *A. lyrata)*. Tetraploids from all four tetraploid lineages of *A. arenosa* and *A. lyrata* (blue) always form a single lineage, suggesting the presence of a single shared tetraploid haplotype at the locus. Further, they form an unresolved network with diploids of multiple species, suggesting diversity of allelic sources of each shared tetraploid haplotypes ('mosaic scenario').

### Supplementary Tables

**Table S1:** Genome-wide nucleotide diversity and Tajima's D of *A. lyrata* populations newly sequenced here, calculated over four-fold degenerate sites.

| pop | ploidy | num_snps | num_sites | num_singletons | nucleotide_diversity | Tajima's_D |
|---|---|---|---|---|---|---|
| VLH | 2 | 110098 | 3443207 | 37098 | 0.013 | 0.10 |
| OSL | 2 | 93965 | 3419890 | 26254 | 0.011 | 0.29 |
| STD | 2 | 108857 | 3823332 | 30475 | 0.012 | 0.30 |
| JOH | 4 | 215901 | 3861780 | 49715 | 0.018 | 0.27 |
| MOD | 4 | 222924 | 3815757 | 57494 | 0.018 | 0.09 |
| PEK | 4 | 166466 | 3663712 | 34317 | 0.015 | 0.34 |
| TEM | 4 | 183885 | 3867146 | 41646 | 0.015 | 0.27 |
| SCT | 4 | 161092 | 3881388 | 34141 | 0.014 | 0.38 |

*Notes: num_snps: number of SNPs in the analysis, num_sites: number of genomic sites considered in the analysis. Nucleotide diversity and Tajima's D for A. arenosa populations are reported in (Monnahan et al. 2019).*

**Table S2: Summary of t**he 17 candidate tetraploid positively selected genes.

| Method | A. lyrata code | A. thaliana code | name | TAIR annotation | N lineages candidate SNPs | N lineages PicMin | PicMin FDR corrected p-value |
|---|---|---|---|---|---|---|---|
| candidate SNPs | AL1G26770 | AT1G14750.1 | SDS | Encodes a meiotic cyclin-like protein, distinct from all other known Arabidopsis cyclins. It is not required for meiotic DSB formation but is necessary for meiotic DSB repair via the homologous chromosome. | 4 | | |
| PicMin + candidate SNPs | AL1G27690 | AT1G15570.1 | CYCA2;3 | A2-type cyclin. Negatively regulates endocycles and acts as a key regulator of ploidy levels in Arabidopsis endoreduplication. Interacts physically with CDKA;1. Expressed preferentially in trichomes and young developing tissues. | 4 | 4 | 0.00170 |
| PicMin + candidate SNPs | AL1G35730 | AT1G22275.1 | ZYP1b | One of two nearly identical proteins (ZYP1a) identified by similarity to transverse filament (TF) proteins. These proteins are involved in chromosome synapsis during meiosis I and localize to the synaptonemal complex (SC). Single mutants have reduced fertility and double mutants (induced by RNAi) have severely reduced fertility. | 4 | 4 | 0.00170 |
| PicMin + candidate SNPs | AL1G56960 | AT1G49590.1 | ZOP1 | Encodes a novel nucleic acid-binding protein that is required for both RdDM (RNA-directed DNA methylation) and pre-mRNA splicing. | 4 | 4 | 0.01552 |
| candidate SNPs | AL1G62040 | AT1G53490.1 | HEI10 | Encodes HEI10, a RING finger-containing protein. HEI10 belongs to a group of proteins well conserved among species known as ZMM. Required for class I crossover formation. The mRNA is cell-to-cell mobile. | 4 | | |
| PicMin + candidate SNPs | AL2G25920 | AT1G67370.1 | ASY1 | meiotic asynaptic mutant 1 (ASY1). ASY1 protein is initially distributed as numerous foci throughout the chromatin. During early G2, the foci are juxtaposed to the nascent chromosome axes to form a continuous axis associated signal. | 4 | 4 | 0.00170 |
| PicMin + candidate SNPs | AL2G37810 | AT1G77600.2 | PDS5b | One of 5 PO76/PDS5 cohesion cofactor orthologs of Arabidopsis. | 4 | 3 | 0.05756 |
| PicMin + candidate SNPs | AL3G24370 | AT3G12690.2 | AGC1.5 | Encodes a putative serine/threonine kinase It is expressed specifically in pollen and appears to function redundantly with AGC1.7 to regulate polarized growth of pollen tubes. | 3 | 3 | 0.04452 |
| PicMin + candidate SNPs | AL3G29960 | AT3G16980.1 | NRPB9A | One of two highly similar, non-catalytic subunits common to nuclear DNA-directed RNA polymerases II, IV and V; homologous to budding yeast RPB9. Appears to be redundant with At4g16265 | 3 | 3 | 0.01509 |
| PicMin + candidate | AL4G29630 | AT2G33845.1 | AT2G33845 | mRNA-binding, OB-fold-like protein, cell to cell mobile | 4 | 4 | 0.00226 |

| | | | | | | | |
|---|---|---|---|---|---|---|---|
| SNPs | | | | | | | |
| PicMin + candidate SNPs | AL4G46460 | AT2G46980.2 | ASY3 | Encodes ASY3, a coiled-coil domain protein that is required for normal meiosis. | 3 | 3 | 0.00255 |
| PicMin + candidate SNPs | AL5G32860 | AT3G52270.1 | TfIIFbeta | Transcription initiation factor IIF, beta subunit; transcription initiation from RNA polymerase II promoter | 4 | 4 | 0.00170 |
| candidate SNPs | AL6G15380 | AT5G05490.2 | SYN1/ REC8 | Encodes a RAD21-like gene essential for meiosis. Encodes a 627 a.a. protein that is slightly longer in the N-terminus than SYN1 BP5. | 2 | | |
| PicMin + candidate SNPs | AL7G13140 | AT4G37630.1 | CYCD5;1 | core cell cycle genes; a quantitative trait gene for endoreduplication. | 4 | 3 | 0.02843 |
| PicMin + candidate SNPs | AL7G35790 | AT4G18490.2 | AT4G18490 | unknown protein; expressed in flower stage 15-18, co-transcription with CYCA2;3 | 4 | 4 | 0.00170 |
| PicMin + candidate SNPs | AL8G25600 | AT5G51340.1 | SCC4 | SCC4 is a tetratricopeptide repeat containing protein and a likely component of a plant cohesion loading complex along with its partner SSC2 It is expressed primarily in dividing cells. Loss of function mutants are embryo lethal, arresting by globular stage. | 3 | 4 | 0.00611 |
| PicMin + candidate SNPs | AL8G44080 | AT5G67260.1 | CYCD3;2 | Encode CYCD3;2, a CYCD3 D-type cyclin. Important for determining cell number in developing lateral organs. Mediating cytokinin effects in apical growth and development. | 3 | 3 | 0.07481 |

**Table S2:** Functional annotation of the 17 tetraploid positively selected genes.

| category | term ID | term description | observed gene count | background gene count | strength | false discovery rate | matching proteins in your network (labels) |
|---|---|---|---|---|---|---|---|
| GO Process | GO:0007049 | Cell cycle | 10 | 489 | 1.52 | 2.96E-10 | SDS,CYCA2;3,ZYP1b,HEI10,ASY1,ASY3,CYCD5;1,SYN1,AT5G51340,CYCD3;2 |
| GO Process | GO:0022402 | Cell cycle process | 9 | 333 | 1.64 | 3.43E-10 | SDS,CYCA2;3,ZYP1b,HEI10,ASY1,CYCD5;1,SYN1,AT5G51340,CYCD3;2 |
| GO Process | GO:0098813 | Nuclear chromosome segregation | 6 | 72 | 2.13 | 8.22E-09 | SDS,ZYP1b,HEI10,ASY1,SYN1,AT5G51340 |
| GO Process | GO:0045143 | Homologous chromosome segregation | 5 | 28 | 2.46 | 1.32E-08 | SDS,ZYP1b,HEI10,ASY1,SYN1 |
| GO Process | GO:0000280 | Nuclear division | 6 | 113 | 1.93 | 5.47E-08 | SDS,ZYP1b,HEI10,ASY1,SYN1,AT5G51340 |
| GO Process | GO:0070192 | Chromosome organization involved in meiotic cell cycle | 5 | 43 | 2.27 | 5.47E-08 | SDS,ZYP1b,HEI10,ASY1,SYN1 |
| GO Process | GO:0051321 | Meiotic cell cycle | 6 | 157 | 1.79 | 2.01E-07 | SDS,ZYP1b,HEI10,ASY1,ASY3,SYN1 |
| GO Process | GO:0007129 | Homologous chromosome pairing at meiosis | 4 | 23 | 2.45 | 6.35E-07 | SDS,ZYP1b,HEI10,ASY1 |
| GO Process | GO:0007131 | Reciprocal meiotic recombination | 4 | 45 | 2.16 | 6.18E-06 | SDS,ZYP1b,HEI10,ASY1 |
| GO Process | GO:0044772 | Mitotic cell cycle phase transition | 4 | 45 | 2.16 | 6.18E-06 | SDS,CYCA2;3,CYCD5;1,CYCD3;2 |
| GO Process | GO:1903047 | Mitotic cell cycle process | 5 | 148 | 1.74 | 6.97E-06 | SDS,CYCA2;3,CYCD5;1,AT5G51340,CYCD3;2 |
| GO Process | GO:0022414 | Reproductive process | 9 | 1429 | 1.01 | 1.08E-05 | SDS,ZYP1b,HEI10,ASY1,ASY3,AGC1.5,SYN1,AT5G51340,CYCD3;2 |
| GO Process | GO:0051301 | Cell division | 6 | 348 | 1.44 | 1.08E-05 | SDS,CYCA2;3,ZYP1b,CYCD5;1,AT5G51340,CYCD3;2 |
| GO Process | GO:0006259 | DNA metabolic process | 6 | 378 | 1.41 | 1.50E-05 | SDS,ZYP1b,HEI10,ASY1,NRPB9A,SYN1 |
| GO Process | GO:0051026 | Chiasma assembly | 3 | 12 | 2.61 | 1.54E-05 | SDS,HEI10,ASY1 |

| | | | | | | | |
|---|---|---|---|---|---|---|---|
| GO Process | GO:0051276 | Chromosome organization | 6 | 487 | 1.3 | 5.43E-05 | SDS,ZYP1b,HEI10,ASY1,SYN1,AT5G51340 |
| GO Process | GO:0051726 | Regulation of cell cycle | 5 | 244 | 1.52 | 5.43E-05 | SDS,CYCA2;3,ASY3,CYCD5;1,CYCD3;2 |
| GO Process | GO:0090304 | Nucleic acid metabolic process | 7 | 1403 | 0.91 | 0.0014 | SDS,ZYP1b,AT1G49590,HEI10,ASY1,NRPB9A,SYN1 |
| GO Process | GO:0010444 | Guard mother cell differentiation | 2 | 11 | 2.47 | 0.0031 | CYCA2;3,CYCD3;2 |
| GO Process | GO:0042023 | DNA endoreduplication | 2 | 19 | 2.23 | 0.0081 | CYCA2;3,CYCD5;1 |
| GO Process | GO:0007062 | Sister chromatid cohesion | 2 | 24 | 2.13 | 0.0119 | SYN1,AT5G51340 |
| GO Process | GO:0044260 | Cellular macromolecule metabolic process | 9 | 3665 | 0.6 | 0.012 | SDS,CYCA2;3,ZYP1b,HEI10,ASY1,AGC1.5,NRPB9A,CYCD5;1,SYN1 |
| GO Process | GO:0043170 | Macromolecule metabolic process | 10 | 4720 | 0.53 | 0.0132 | SDS,CYCA2;3,ZYP1b,AT1G49590,HEI10,ASY1,AGC1.5,NRPB9A,CYCD5;1,SYN1 |
| GO Process | GO:0006281 | DNA repair | 3 | 260 | 1.27 | 0.042 | SDS,NRPB9A,SYN1 |
| GO Function | GO:0016538 | Cyclin-dependent protein serine/threonine kinase regulator activity | 4 | 52 | 2.09 | 7.64E-05 | SDS,CYCA2;3,CYCD5;1,CYCD3;2 |
| GO Component | GO:0000794 | Condensed nuclear chromosome | 4 | 27 | 2.38 | 2.57E-06 | ZYP1b,HEI10,ASY1,SYN1 |
| GO Component | GO:0000307 | Cyclin-dependent protein kinase holoenzyme complex | 4 | 48 | 2.13 | 8.39E-06 | SDS,CYCA2;3,CYCD5;1,CYCD3;2 |
| GO Component | GO:0005694 | Chromosome | 6 | 300 | 1.51 | 8.39E-06 | ZYP1b,HEI10,ASY1,ASY3,SYN1,AT5G51340 |
| GO Component | GO:0061695 | Transferase complex, transferring phosphorus-containing groups | 5 | 169 | 1.68 | 9.24E-06 | SDS,CYCA2;3,NRPB9A,CYCD5;1,CYCD3;2 |
| GO Component | GO:0005634 | Nucleus | 13 | 4669 | 0.65 | 1.26E-05 | SDS,CYCA2;3,ZYP1b,AT1G49590,HEI10,ASY1,ASY3,AGC1.5,NRPB9A,CYCD5;1,SYN1,AT5G51340,CYCD3;2 |
| GO Component | GO:0031981 | Nuclear lumen | 7 | 807 | 1.15 | 2.70E-05 | ZYP1b,AT1G49590,HEI10,ASY1,ASY3,NRPB9A,SYN1 |
| GO Component | GO:0043232 | Intracellular non-membrane-bounded organelle | 7 | 1407 | 0.9 | 0.00064 | ZYP1b,HEI10,ASY1,ASY3,NRPB9A,SYN1,AT5G51340 |
| GO Component | GO:0032991 | Protein-containing complex | 8 | 2226 | 0.76 | 0.0011 | SDS,CYCA2;3,AT1G49590,NRPB9A,CYCD5;1,SYN1,AT5G51340,CYCD3;2 |
| GO Component | GO:0042025 | Host cell nucleus | 3 | 150 | 1.51 | 0.0047 | SDS,CYCA2;3,CYCD3;2 |
| GO Component | GO:0005654 | Nucleoplasm | 3 | 396 | 1.09 | 0.0472 | AT1G49590,ASY1,NRPB9A |
| UniProt Keywords | KW-0469 | Meiosis | 6 | 70 | 2.14 | 2.09E-09 | SDS,ZYP1b,HEI10,ASY1,ASY3,SYN1 |
| UniProt Keywords | KW-0131 | Cell cycle | 7 | 266 | 1.63 | 3.74E-08 | SDS,CYCA2;3,ZYP1b,CYCD5;1,SYN1,AT5G51340,CYCD3;2 |
| UniProt Keywords | KW-0132 | Cell division | 6 | 198 | 1.69 | 2.83E-07 | SDS,CYCA2;3,ZYP1b,CYCD5;1,AT5G51340,CYCD3;2 |
| UniProt Keywords | KW-0195 | Cyclin | 4 | 52 | 2.09 | 4.24E-06 | SDS,CYCA2;3,CYCD5;1,CYCD3;2 |
| UniProt Keywords | KW-0158 | Chromosome | 3 | 117 | 1.62 | 0.005 | HEI10,ASY1,ASY3 |
| UniProt | KW- | Chromosome partition | 2 | 24 | 2.13 | 0.0091 | SYN1,AT5G51340 |

| | | | | | | | |
|---|---|---|---|---|---|---|---|
| Keywords | 0159 | | | | | | |
| UniProt Keywords | KW-0539 | Nucleus | 9 | 3770 | 0.59 | 0.0101 | CYCA2;3,ZYP1b,AT1G49590,HEI10,ASY1,ASY3,NRPB9A,SYN1,AT5G51340 |
| STRING clusters | CL:6994 | Cell division, and microtubule-based movement | 7 | 164 | 1.84 | 3.22E-08 | SDS,CYCA2;3,AT1G77600,CYCD5;1,SYN1,AT5G51340,CYCD3;2 |
| STRING clusters | CL:7443 | Homologous chromosome pairing at meiosis, and meiosis protein spo22/zip4 like | 4 | 13 | 2.7 | 6.45E-07 | ZYP1b,HEI10,ASY1,ASY3 |
| STRING clusters | CL:7001 | Cyclin, and RNA polymerase II CTD heptapeptide repeat kinase activity | 4 | 38 | 2.23 | 1.20E-05 | SDS,CYCA2;3,CYCD5;1,CYCD3;2 |
| STRING clusters | CL:7270 | Cohesin complex, and smc loading complex | 3 | 10 | 2.68 | 4.34E-05 | AT1G77600,SYN1,AT5G51340 |
| STRING clusters | CL:7005 | Cyclins are a family of proteins that control the progression of cells through the cell cycle by activating cyclin-dependent kinase (Cdk) enzymes., and Cyclin-dependent kinase regulatory subunit | 3 | 23 | 2.32 | 0.00027 | CYCA2;3,CYCD5;1,CYCD3;2 |
| STRING clusters | CL:7007 | Cyclins are a family of proteins that control the progression of cells through the cell cycle by activating cyclin-dependent kinase (Cdk) enzymes., and regulation of stomatal complex patterning | 2 | 12 | 2.43 | 0.0126 | CYCA2;3,CYCD3;2 |

**Table S4:** Presence and frequency of tetraploid, two diploid, and other haplotype blocks in all 61 tetraploid populations of *A. arenosa* and *A. lyrata* (479 individuals). AF: allele frequency of haplotype.

| Gene | AF (tetraploid) | AF (*A. arenosa* diploid) | AF (*A. lyrata* diploid) | other | Tetraploid haplotype in N of pops | total N of tetraploid pops | % of presence |
|---|---|---|---|---|---|---|---|
| CYCA2_3 | 0.47 | 0.02 | 0.04 | 0.48 | 56 | 61 | 91.8% |
| CYCD3_2 | 0.61 | 0.13 | 0.06 | 0.20 | 59 | 60 | 98.3% |
| CYCD5_1 | 0.52 | 0.12 | 0.07 | 0.30 | 56 | 60 | 93.3% |
| AT4G18490 | 0.57 | 0.10 | 0.04 | 0.30 | 57 | 60 | 95.0% |
| ZYP1b | 0.68 | 0.05 | 0.08 | 0.18 | 57 | 58 | 98.3% |
| ASY1 | 0.62 | 0.06 | 0.02 | 0.31 | 57 | 60 | 95.0% |
| PDS5b | 0.73 | 0.02 | 0.02 | 0.23 | 58 | 60 | 96.7% |
| ASY3 | 0.61 | 0.01 | 0.01 | 0.36 | 56 | 59 | 94.9% |
| SCC4 | 0.57 | 0.16 | 0.03 | 0.23 | 59 | 61 | 96.7% |
| NRPB9A | 0.73 | 0.06 | 0.02 | 0.19 | 47 | 50 | 94.0% |
| TfIIFbeta | 0.85 | 0.03 | 0.01 | 0.12 | 55 | 58 | 94.8% |
| AT2G33845 | 0.51 | 0.04 | 0.02 | 0.42 | 58 | 60 | 96.7% |

#### Supplementary Datasets

Dataset S1: Metadata and sequence quality checks for the 983 whole genome sequenced individuals.

Dataset S2: Set of 54 significant PicMin windows.

Dataset S3: List of outlier genes identified using the 'candidate SNP' approach and their overlap among tetraploid lineages.

Dataset S4: Table of linked candidate SNPs, as determined using long read sequencing. Position of variants within the same column (columns *D-N*) highlights their physical linkage within a gene region (black boxes). Note that these linked candidate SNPs were used as markers to reconstruct haplotypes from short read data, which is summarised in columns *Q-AND*.

Dataset S5: Sequences of the 12 positively selected genes (PSGs), assembled using long read sequencing of five diploid and five tetraploid individuals.

Dataset S6: Candidate SNPs marking haplotypes of CYCA2;3 and PDS5b (Fig. 4C, D), as found on the same long read.

Dataset S7: Distribution of the 232 tetraploid candidate SNPs among the 504 diploid samples used to estimate the likely sources of tetraploid haplotypes.

Dataset S8: Sampling locations of the 129 populations.
